# Planar cell polarity signaling controls cell division symmetry to promote termination of adult tissue regeneration

**DOI:** 10.1101/2025.04.27.650884

**Authors:** Samantha J. Hack, Wendy S. Beane

**Affiliations:** Department of Biological Sciences, Western Michigan University, Kalamazoo, MI 49008, USA; Department of Cell and Molecular Biology, St. Jude Children’s Research Hospital, Memphis, TN 38105, USA

**Keywords:** Regeneration, planar cell polarity, growth termination, stem cells, proliferation, allometry

## Abstract

Tissue formation is coordinated by cell-intrinsic and cell-extrinsic signals across space and time, yet how self-limiting growth is controlled remains mysterious. Here, we leveraged the highly regenerative planarian *Schmidtea mediterranea* to identify molecular regulators of endogenous growth termination in adults, where unchecked growth can promote carcinogenesis. We identified the Planar Cell Polarity (PCP) pathway as a key regulator of body-wide regenerative growth through control of stem cell division symmetry. Following PCP pathway inhibition, widespread tissue hyperplasia occurred weeks after regeneration normally finishes. Importantly, this was due to progenitor expansion and depletion of adult *tspan-1^+^* pluripotent stem cells capable of whole-body regeneration. Using transcriptional analysis of regenerating animals over time, we identified that control of stem cell fate by changes in cell division symmetry is a potential growth termination mechanism. While symmetric divisions maintain the stem cell pool in adult planaria, upon PCP loss, the number of asymmetrically dividing cells increases, driving stem cell depletion and excessive tissue differentiation. Our data suggest that PCP signaling regulates stem cell maintenance and fate decisions during self-limiting growth.

**Summary Statement:** Here, we show that Planar Cell Polarity (PCP) signaling regulates stem cell division symmetry to control termination of adult tissue regeneration in planarians.

## Introduction

Regeneration is driven by mechanisms that identify lost or damaged cell types and programs that integrate new cells into existing structures to maintain overall tissue pattern, size, and function. Importantly, successful completion of self-limiting regeneration relies not only on the regulation and coordination of stem cell proliferation and differentiation to form new structures, but also on the appropriate repression of these processes in time and space to prevent hyperplastic growth (<u>Marinari et al., 2012</u>; <u>Takemura and Nakato, 2017</u>). However, how growth termination cues are integrated with patterning information and how they instruct stem cell function to repress growth remains unclear.

Most of what is currently known about the termination phase of regeneration has been uncovered by studies of the mammalian liver, a stunning example of self-limiting growth *in situ*. Removal of up to two thirds of the murine liver by partial hepatectomy results in hepatocyte proliferation and hypertrophy that restores the original liver mass within 3 weeks (<u>Michalopoulos and Bhushan, 2021</u>). While hepatocyte proliferation is known to be limited to the first week post injury [followed by upregulation of differentiation transcriptional programs, increased ECM deposition, and the return of proliferating cells to a quiescent state (<u>Donthamsetty et al., 2013</u>; <u>Gallai et al., 1996</u>; <u>Michalopoulos and Bhushan, 2021</u>; <u>Rudolph et al., 1999</u>)], the timing of growth termination is unclear. During embryogenesis, self-organization is instructed by cell-intrinsic genetic and molecular programs that cooperate to prevent hyperplastic growth. Signal transduction pathways, and in particular the Hippo pathway (and its transcriptional regulator Yorkie; YAP and TAZ) and the mTOR pathway, have been implicated in the regulation of overall organ size and growth regulation (<u>Csibi and Blenis, 2012</u>; <u>Tumaneng et al., 2012</u>; <u>Yu et al., 2015</u>). While such embryonic regulators represent enticing candidates for controllers of adult tissue morphogenesis, the extent to which these endogenous growth-termination mechanisms are recapitulated during regeneration and whether they can be manipulated to limit new growth *in situ* remains to be addressed. One key difference between developmental and adult morphogenesis is that adult tissue growth begins from a pre-existing pattern and form. Thus, regulators of overall tissue pattern that also control stem cell states and proliferative capacity, such as the planar cell polarity (PCP) pathway, are logical candidates for potential growth termination signals.

The PCP pathway is a highly conserved non-canonical Wnt pathway classically associated with controlling cell polarization. Specifically, it regulates asymmetric localization of the protein complexes Vangl/Prickle and Frizzled/Dishevelled/Diego to coordinate proximal-distal directionality of cells across a planar tissue (<u>Butler and Wallingford, 2017</u>; <u>Struhl et al., 2012</u>; <u>Strutt et al., 2011</u>). The cell-cell communication that stabilizes the PCP pattern and propagates polarity information across distant cells to entire tissues is mediated by homodimers formed between the extracellular domains of the atypical cadherin protein Flamingo (Fmi/CELSR) (<u>Chen et al., 2008</u>; <u>Usui et al., 1999</u>). In addition, core PCP proteins govern regulation of cytoskeletal organization and actomyosin dynamics (<u>Kim et al., 2010</u>; <u>Shindo and Wallingford, 2014</u>; <u>Vladar et al., 2012</u>; <u>Winter et al., 2001</u>), for example in cell division orientation, cell cycle regulation, and maintenance of stem cell quiescence (<u>Chavali et al., 2018</u>; <u>Ju et al., 2010</u>; <u>Ségalen et al., 2010</u>; <u>Yang et al., 2017</u>). Moreover, as PCP proteins undergo endosomal recycling during mitosis and are inherited by daughter cells, polarity information is retained during proliferation, suggesting that PCP signaling plays an important role in directing stem cell activities during regenerative morphogenesis (<u>Devenport et al., 2011</u>; <u>Heck and Devenport, 2017</u>; <u>Shrestha et al., 2015</u>). This is consistent with studies suggesting that PCP signaling regulates stem cell proliferation during adult tissue regeneration. For example, Vangl2 is required to reorient the mitotic spindle of proliferating neural stem cells to correctly direct morphogenesis of the regenerating axolotl spinal cord (<u>Rodrigo Albors et al., 2015</u>).

Planarian flatworms notoriously undergo robust whole-body regeneration from small tissue fragments (Reddien, 2018) while avoiding hyperplastic growth and tumorigenesis (<u>Barghouth et al., 2019</u>). Thus, planaria are a unique model to uncover the basic biological mechanisms of endogenous growth termination. In previous work, we reported that loss of PCP signaling in the planarian *Schmidtea mediterranea* and in *Xenopus* tadpoles results in neural hyperplasia and stem cell hyperproliferation that continues weeks after normal regeneration has ceased (<u>Beane et al., 2012a</u>). Here, we show that inhibition of PCP signaling in planarians results in body-wide tissue hyperplasia due to dysregulation of stem cell division symmetry and differentiation dynamics. Hyperproliferation resulted in expanded progenitor populations and depletion of pluripotent cells, suggesting that PCP signaling may be required to maintain stem cell quiescence during growth termination and tissue maintenance. Interestingly, while in controls the proportion of symmetrically dividing stem cells increased over time (expanding the tissue maintenance stem cell pool), levels of asymmetric cell division and continued progenitor cell formation remained high in PCP-inhibited animals. Collectively, our data shows that PCP signaling is required to support the maintenance of pluripotent stem cell populations and ensure self-limitation of new growth during adult tissue regeneration.

## Results

*Transcriptome-Wide Analysis of Regenerating Planaria Upon Inhibition of PCP Signaling* We previously showed that ectopic visual structures and commissural neuronal connections in the planarian central nervous system (CNS) increased in abundance over time upon loss of PCP signaling (<u>Beane et al., 2012a</u>). This phenotype suggests that PCP signaling may be required to restrict regrowth in at least two cell lineages, since photoreceptor neurons and brain commissural neurons arise from separate progenitor populations (<u>Cowles et al., 2013</u>; <u>Cowles et al., 2014</u>; <u>Lapan and Reddien, 2011</u>; <u>Lapan and Reddien, 2012</u>). To identify tissue types that may require PCP signaling for growth termination and to uncover potential growth termination genes, we performed bulk RNAseq analysis of control and PCP-inhibited planarians (*Schmidtea mediterranea*) undergoing whole body regeneration (2-weeks) and homeostatic tissue maintenance post regeneration (8-weeks) (Fig. 1A). RNA interference (PCP RNAi) with dsRNA targeting Vangl2 (*smed-vang-1*), Rho-associated coiled-coil protein kinase (ROCK, *smed-rock*), and dishevelled associated activator of morphogenesis (DAAM, *smed-daam-1*) was conducted as previously described (<u>Beane et al., 2012b</u>) (Fig. 1B-D). After treatment, planarians were amputated into trunk fragments and allowed to regenerate for two or eight weeks before total RNA extraction and bulk mRNA sequencing (Fig. 1A, Table S1). To identify transcriptional changes across conditions, we conducted differential expression analysis where all genes with -0.5 ≤ log_2_fold change ≥ 0.5 and *p-adj*. < 0.05 were deemed to be significantly differentially expressed (DEGs) (Fig. 1E, Table S2).

**Figure 1:**
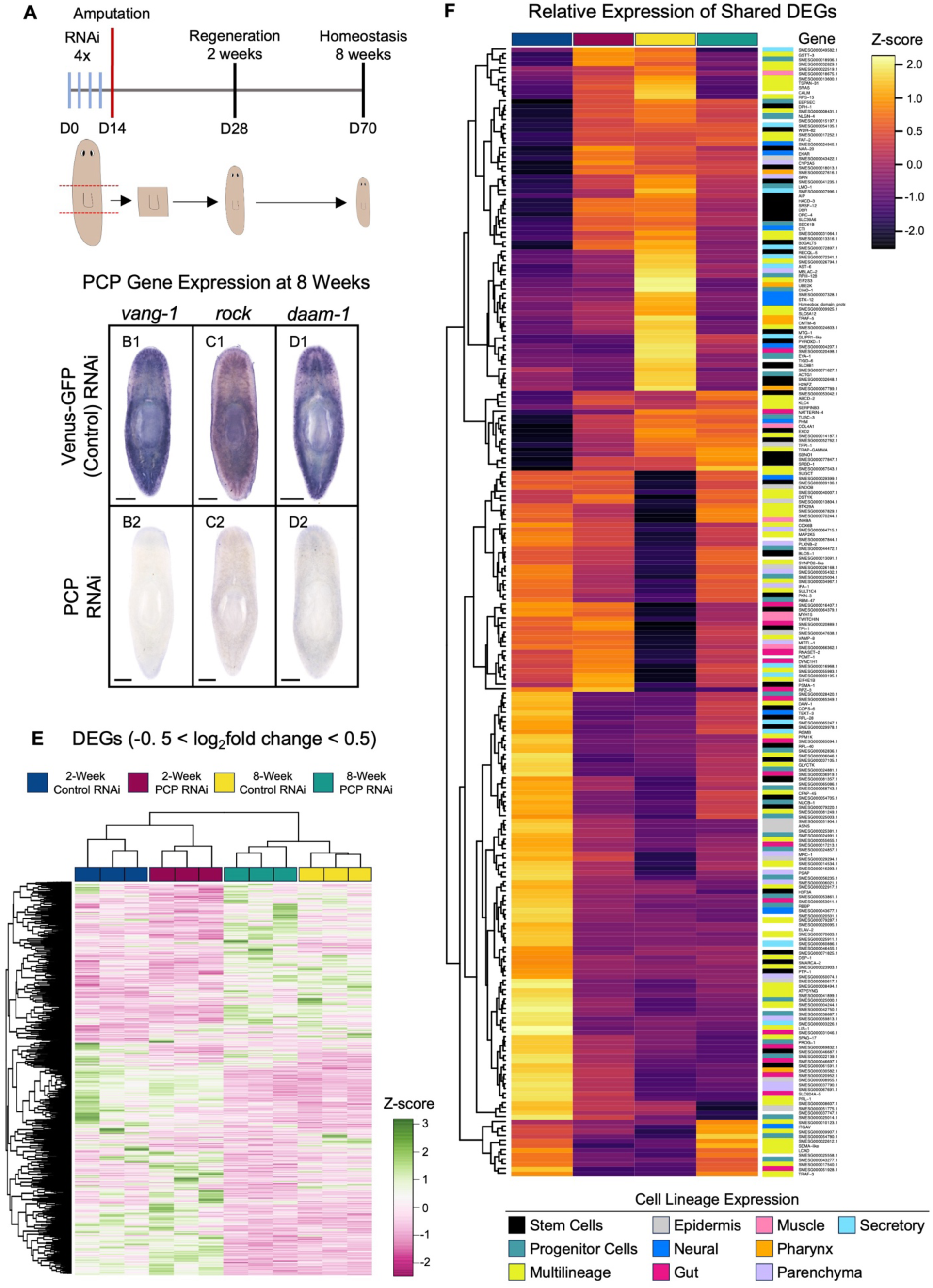
Bulk RNA-seq identifies potential PCP-mediated growth termination genes. **(A)** Experimental timeline. **(B-D)** Expression of *vang-1*, *rock*, or *daam-1* at 8 weeks in Control and PCP RNAi animals. Scale bars: 250 µm. n ≥ 7. **(E)** Significantly differentially expressed genes (DEGs, -0.5 ≥ log_2_fold change ≥ 0.5, *p-adj.* < 0.05) between Control and PCP-inhibited animals at 2-weeks and 8-weeks. **(F)** Relative expression (Z-score of average FPKM) of 240 shared genes in groups of interest from (Fig. S1B). The major cell lineage type for each gene is displayed to the right. All experiments were repeated in triplicate.

Genes that did not fall into the log_2_fold change cutoff but had *p-adj.* < 0.05 are also included in Table S2. We used normal hierarchical clustering to visualize the relative expression of all DEGs (Fig. 1E). Based on clustering analysis, PCP RNAi conditions had more similar expression patterns with their time-matched controls than between PCP-inhibited animals at 2- and 8-weeks. Similarly, whole transcriptome principal component analysis (PCA) showed that most variance occurred between timepoints (Fig. S1A). Together, these data suggest that large transcriptional differences exist between animals at the end stages of regeneration (2-weeks) and those undergoing homeostatic tissue maintenance (8-weeks).

Next, we cross-compared all DEGs (*p-adj.* < 0.05) between conditions to select for groups of genes for which expression was affected by PCP RNAi at both timepoints (Fig. S1B). We did not use a log_2_fold change cutoff for this analysis in order to avoid excluding genes that may have different expression levels across conditions. Genes identified in all comparisons in Fig. S1B are listed in Table S3. For all genes of interest (Fig. S1B), we identified the homologous transcript in the dd_Smed_v6 transcriptome (<u>Grohme et al., 2018</u>; <u>Rozanski et al., 2019</u>) and distinguished its major cell lineage expression in the Planarian Single Cell Atlas (<u>Plass et al., 2018</u>) (Fig. 1F). Coupled with complete hierarchical clustering of the relative expression (Z-score FPKM) of each gene, this analysis allowed us to visualize trends in expression and tissue lineages that may require PCP signaling for regeneration, maintenance, or growth termination.

Most of the shared genes between conditions had enriched expression in the main cell type lineages including stem cells, stem cell progeny, the gut (goblet cells, phagocytes, and gut progenitors), secretory cells, and the epidermis (epidermis, dorsal-ventral boundary, and epidermal progenitors) (Fig. 1F). Other shared DEGs had diverse, multilineage expression in one or more major cell types including neural, pharynx, protonephridia, secretory, parenchymal, muscle, and gut lineages (Fig. 1F). Indeed, *vang-1*, *rock*, and *daam-1* are also expressed in multiple tissue lineages; *vang-1* expression is highly enriched in neurons (GABA and ChAT), muscle, and activated early epidermal progenitors while *rock* has enriched expression in neurons and the epidermis, in addition to high expression in the pharynx, gut, and secretory cells. Finally, *daam-1* expression is enriched in almost every major differentiated cell lineage including pigment cells, neurons, the gut, parenchymal cells, protonephridia, glia, and the epidermis (<u>Plass et al., 2018</u>). Thus, it is not surprising that transcriptional changes in genes expressed in almost all cell type lineages were detected following PCP RNAi (Fig. 1F). Collectively, our differential expression analyses indicate that dynamic and unique gene expression changes occur between the end stages of regeneration and the initiation of tissue maintenance programs. Furthermore, these data show that PCP RNAi causes transcriptional changes in genes expressed in every major cell-type lineage, supporting our hypothesis that PCP signaling serves as a body-wide growth termination signal.

### Loss of PCP Signaling Results in Multi-Organ System Tissue Hyperplasia

Next, we asked whether PCP signaling may be required for terminating growth throughout the body. To answer this question, we analyzed expression of differentiated tissue markers and evaluated effects on gross morphology in multiple organ systems at 8-weeks in control and PCP-inhibited animals, starting with the intestinal system. The planarian gut is composed of three main cell types (secretory goblet cells, absorptive phagocytes, and basal cells) (<u>Fincher et al., 2018</u>; <u>Forsthoefel et al., 2020</u>). Its highly branched intestine has been proposed to serve as a potential niche for adult stem cells (<u>Barberan et al., 2016</u>; <u>Flores et al., 2016</u>; <u>Forsthoefel et al., 2012</u>; <u>Henderson et al., 2015</u>). The gut has one anterior primary intestinal branch and two posterior primary branches, with secondary branches projecting laterally and forming tertiary (and sometimes quaternary) branches that extend to the margins of the animal (<u>Forsthoefel et al., 2012</u>; <u>Forsthoefel et al., 2011</u>) (Fig. 2A). To quantify effects on gut branch numbers, as well as the dimensions of individual secondary branches at 8-weeks following PCP inhibition, we visualized intestinal branching morphology by whole-mount colorimetric *in situ* hybridization (WISH) using the intestinal phagocyte marker *innexin-9* (*inx-9*) (Fig. 2B, D, F, Fig. S2A). We normalized the total number of secondary branches to animal size and quantified the number of tertiary branches for each secondary branch. We found that the average number of secondary branches per mm^2^ of body area was not affected in either the anterior or posterior gut following PCP loss (p ≥ 0.05, Fig. 2C). Similarly, control and PCP-inhibited animals had comparable numbers of tertiary branches (p ≥ 0.05, Fig. S2B).

**Figure 2:**
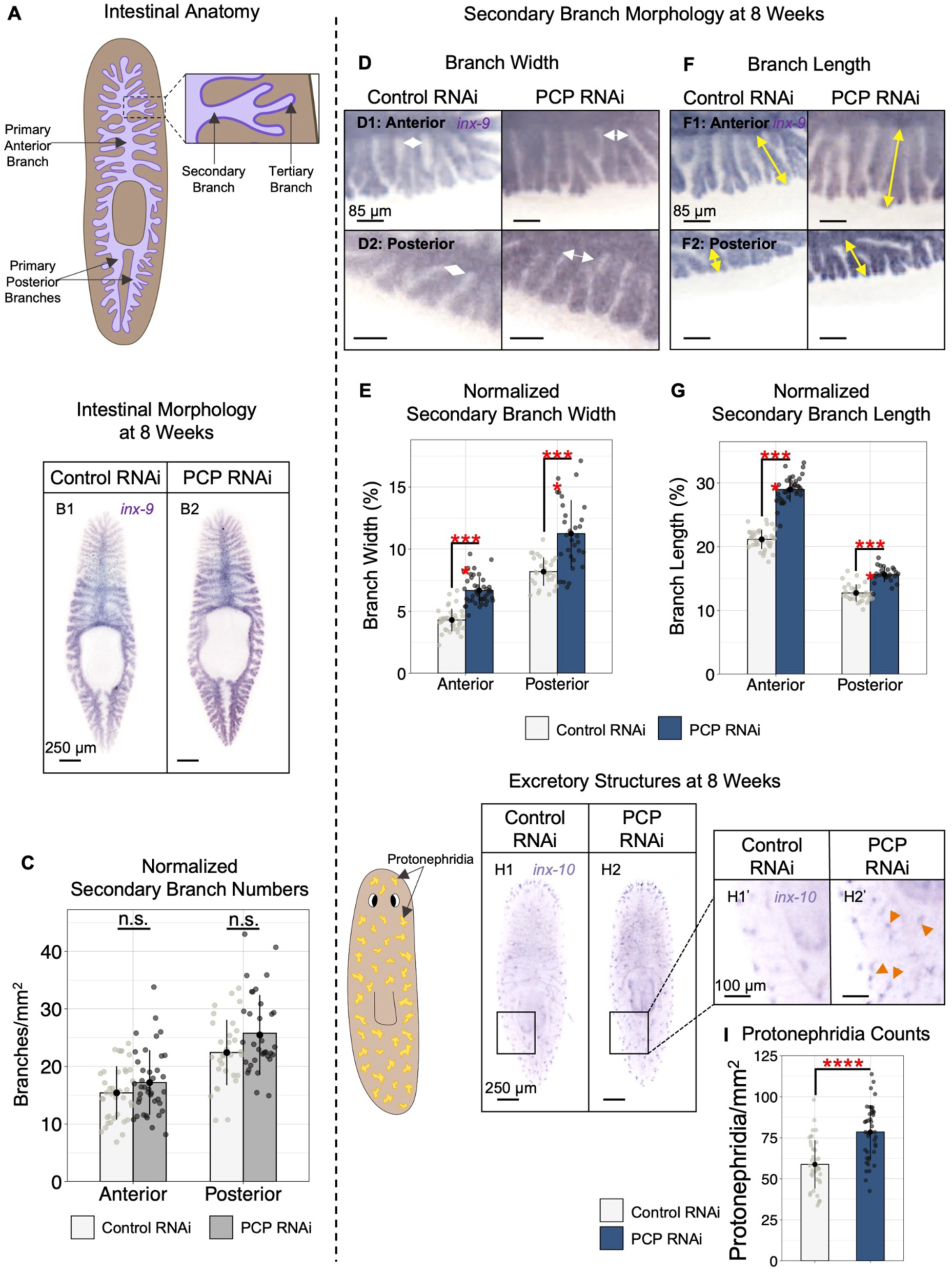
Inhibition of PCP signaling causes multi-organ system hyperplasia following regeneration. **(A)** Planarian intestinal anatomy. **(B)** Intestinal morphology visualized by *inx-9* expression at 8 weeks. Scale bars: 250 µm. Anterior: up. **(C)** Quantification of (B); Mean±s.d. secondary intestinal branch numbers at 8 weeks. n ≥ 32. Two sample *t*-test: no significant difference (n.s.). **(D-G)** Morphology of anterior (D1, F1) and posterior (D2, F2) secondary branches. Scale bars: 85 µm. White double arrowheads: Branch width. Yellow double arrowheads: Branch length. **(E)** Quantification of (D); Mean±s.d. normalized secondary branch width. n ≥ 31. Two sample *t*-test: **** p ≤ 0.0001. **(G)** Quantification of (F); Mean±s.d. normalized secondary branch length. n ≥ 25. Two sample *t*-test: **** p ≤ 0.0001. **(H)** Protonephridia morphology at 8 weeks visualized by *inx-10* expression. Scale bars: 250 µm. Anterior: up. H1’ and H2’: Zoom of region in H1 and H2. Orange arrows: ectopic excretory structures. Scale bars: 100 µm. **(I)** Quantification of (H); Mean±s.d. protonephridia numbers at 8 weeks (protonephridia/mm^2^ body area). n ≥ 41. Two sample *t*-test: **** p ≤ 0.0001. All experiments were repeated in triplicate.

In an earlier study, the intestine was found to grow by increasing the size of primary and secondary branches, in addition to increasing the numbers of secondary and tertiary branches (<u>Forsthoefel et al., 2011</u>). This analysis implies that the number of gut branches may not reflect the total intestinal tissue within an animal at any given time. Thus, we next compared the dimensions of individual secondary branches at 8-weeks following PCP loss with those of secondary branches in control RNAi animals (Fig. 2D-G). In this analysis, we found that PCP loss led to significant increases in both secondary branch width and length (p ≤ 0.0001 for both, Fig. 2E, G). Taken together, these data indicate that PCP signaling is required to restrict intestinal regrowth, consistent with our previous findings in neural tissue (<u>Beane et al., 2012a</u>).

In addition to enriched expression in gut cell types like phagocytes, PCP gene expression is also enriched in protonephridia and their progenitors (<u>Plass et al., 2018</u>). Protonephridia are ciliated excretory structures analogous to the vertebrate kidney that are capable of forming cyst-like malformations in their tubular assemblies (<u>Thi-Kim Vu et al., 2015</u>). Considering that loss of PCP signaling is associated with formation of ciliopathies, such as polycystic kidney disease, and coupled with the enriched expression in protonephridia, we asked whether PCP signaling might also regulate the morphogenesis of excretory structures. Using WISH for *innexin-10* (*inx-10*), we visualized the distribution of protonephridia at 8-weeks in control and PCP RNAi animals (Fig. 2H). When normalized to body area, we found that the density of excretory structures was significantly increased following PCP-inhibition (p ≤ 0.0001, Fig. 2I), concomitant with protonephridia that were slightly smaller in overall size. Finally, we assessed changes in morphology of the pharynx, a muscular tube with an extensive neural network responsible for nutrient intake and waste removal (<u>Miyamoto et al., 2020</u>). When combining nuclei labeling with detection of *laminin* expression by fluorescent *in situ* hybridization (FISH), the pharynx is easily visualized on the ventral side of the animal (<u>Adler et al., 2014</u>). Relative to body size at 8-weeks (Fig. S2C,D), we found that pharynges in PCP-inhibited animals were on average 26% larger relative to controls (p ≤ 0.01, Fig. S2E). Collectively, our morphological analyses indicate that PCP signaling is required for termination of tissue regeneration in multiple organ systems in planarians, and that loss of PCP signaling leads to body-wide tissue hyperplasia.

### Regulation of Cell Number and Size is Altered Following PCP Inhibition

In vertebrates, PCP signaling is a well-known regulator of epidermal cell polarity and controls orientation of cell division (<u>Devenport et al., 2011</u>; <u>Luxenburg et al., 2015</u>; <u>Oozeer et al., 2017</u>). Similarly in planarians, PCP is required for organism-wide organization of ventral epidermal cilia (<u>Almuedo-Castillo et al., 2011</u>; <u>Hanh Thi-Kim et al., 2019</u>). To investigate how PCP RNAi affects epidermal morphology, we quantified cell nuclei and the filamentous actin (F-actin) network of the epidermis in control and PCP RNAi animals at 2- and 8-weeks in a 50 mm^2^ area anterior to the pharynx (white boxed region on graphic to the left, Fig. 3A-B). Total epidermal cell numbers were quantified within one area to account for potential changes in cell density in differing anatomical regions. At 2 weeks, when regenerative processes are still occurring, epidermal cell density did not differ between control and PCP-inhibited animals (p ≥ 0.05, Fig. 3A,D) and the F-actin network had similar morphology between conditions (Fig. 3A’’). At 8-weeks (which is more than 4 weeks after regeneration is normally considered finished), epidermal cell density was significantly increased in PCP-inhibited animals (p ≤ 0.0001, Fig. 3B,D). In addition to the observed increase in epidermal cell nuclei, we also noted that F-actin deposition appeared to be increased following PCP loss (Fig. 3B’’).

**Figure 3:**
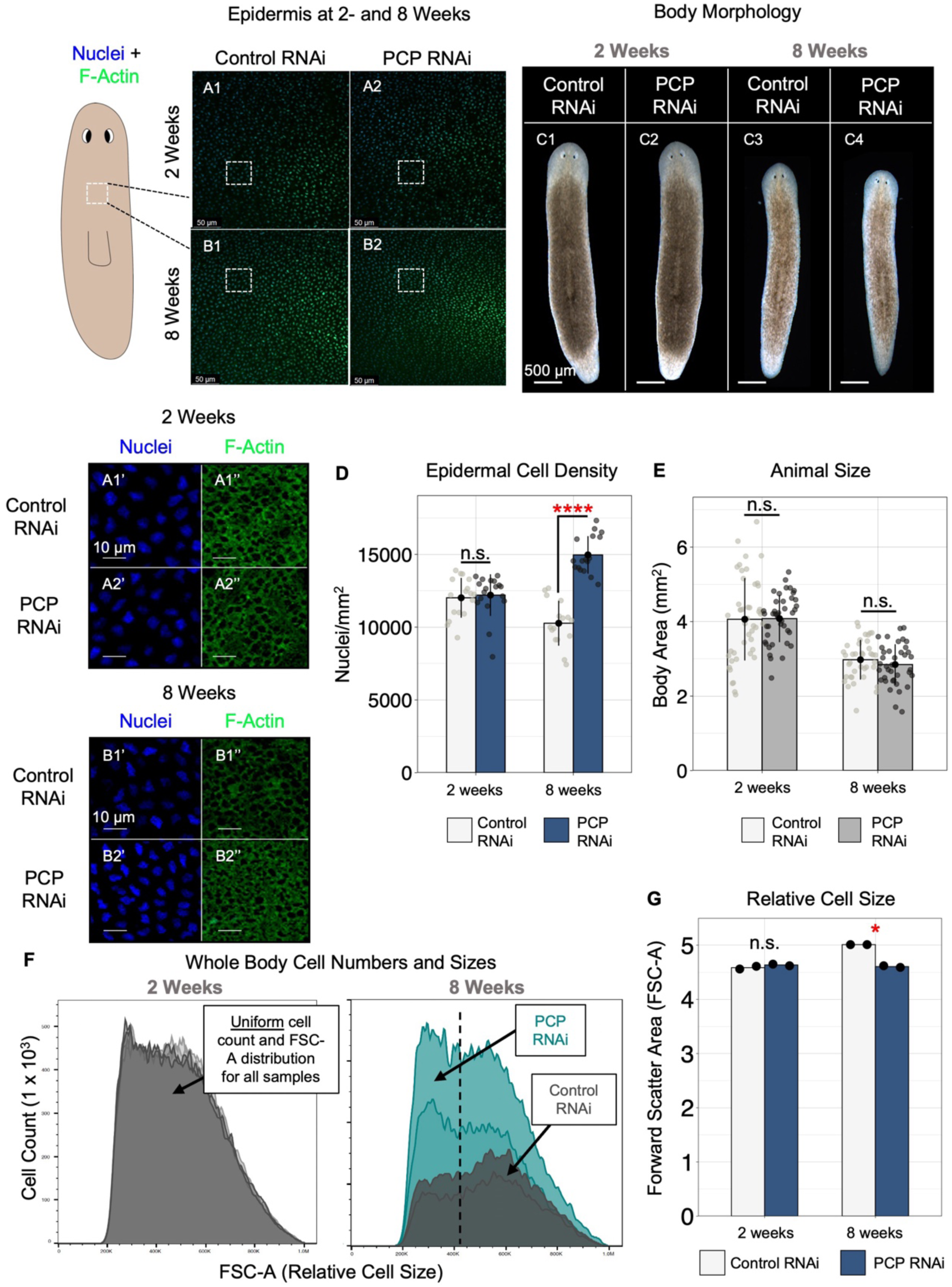
Epidermal cell density increases following PCP inhibition. **(A)** Epidermis in the trunk at 2- and 8-weeks following PCP inhibition. Scale bars: 50 µm. Nuclei (A1’-B2’) and F-actin (A1’’-B2’’) are shown separately for regions in white boxes (A1-B2). Scale bars for all: 10 µm. **(C)** Body morphology at 2-(C1-C2) and 8-(C3-C4) weeks post amputation. Scale bars: 500 µm. Anterior is up. **(D)** Quantification of (A-B); Mean±s.d. epidermal cell density (cells/mm^2^) at 2- and 8 weeks. n ≥ 16. Two sample *t*-test: n.s. (2 weeks) and **** p ≤ 0.0001 (8 weeks). **(E)** Quantification of (C); Mean±s.d. animal size (body area in mm^2^) at 2 and 8 weeks. n ≥ 38. Two sample *t*-test: n.s. for controls compared to PCP RNAi, **** p ≤ 0.0001 Control RNAi at 2 and 8 weeks and PCP at 2 and 8 weeks. **(F)** Flow cytometric analyses of pooled whole-animal lysates at 2- and 8 weeks. Total cell numbers and forward scatter area (FSC-A) for all singlets identified after gating are shown. For each condition at 2- and 8-weeks: 2 pooled replicates of n = 33 animals. **(G)** Quantification of (F): Relative cell size (FSC-A) of all singlets in each condition. Two sample *t*-test: * p < 0.05. Data shown in A-E; repeated in triplicate. Data shown in F-G: Repeated in duplicate.

Given the hyperplastic phenotypes in the gut, excretory system, pharynx, and epidermis, where cell numbers appeared to increase across tissues, we wondered whether PCP inhibition may lead to changes in overall animal size. To address this question, we live-imaged control and PCP-inhibited animals at 2- and 8-weeks and compared total body area (mm^2^) between conditions. Consistent with animals undergoing the processes of degrowth (<u>González-Estévez, 2009</u>; <u>Gonzalez-Estevez et al., 2012</u>), we observed a significant decrease in animal size in both control and PCP-inhibited animals from 2- to 8-weeks (p ≤ 0.0001 for both, Fig. 3C,E). However, no significant difference in body size between control and PCP-inhibited animals was detected at either timepoint (p ≥ 0.05 for both, Fig. 3C,E). Thus, while PCP loss leads to increased cell density, these data suggest that overall animal size is not controlled by PCP signaling.

We hypothesized that cell sizes may be reduced to account for increasing cell numbers to maintain overall body size after PCP inhibition. To investigate changes in whole body cell numbers and sizes, we analyzed whole animal lysates at both 2- and 8 weeks by flow cytometry (Fig. 3F-G). Consistent with our analysis of epidermal cell numbers, PCP inhibition did not affect whole body cell numbers at 2-weeks, but did appear to increase whole-body cell numbers at 8-weeks (Fig. 3F). We then analyzed changes in forward scatter area (FSC-A; a metric of relative cell size), at both 2- and 8-weeks in each condition (Fig. 3G). Similar to cell numbers, relative cell sizes (Fig. 3G) were very similar between control and PCP-inhibited animals at 2-weeks (close overlap of grey distributions on the left of Fig. 3F). FSC-A had a left-shifted distribution in suspensions from PCP-inhibited animals at 8-weeks, which is indicative of smaller relative cell sizes (dashed line in right panel of Fig. 3F shows shift in FSC-A, Fig. 3G; p < 0.05). Together, these data show that while animals degrow in body size over time, counterintuitively, relative total body cell numbers increase after PCP loss, a contradiction that can be explained by reduced cell sizes.

### Neuronal Composition Fails to Scale Allometrically with Brain Size After PCP Loss

Given these observations, we wondered whether increased cell density could be observed in other differentiated tissue lineages. The planarian brain is an excellent model to investigate this question since each neuronal subtype maintains a constant ratio relative to brain size (<u>Takeda et al., 2009</u>), has distinct regional organization (Fig. 4A), and the lineages that arise from different neural progenitor populations are well characterized (<u>Cowles et al., 2013</u>; <u>Cowles et al., 2014</u>; <u>Ross et al., 2017</u>). We assessed expression of prohormone convertase-2 (*smed-pc2*), a neuropeptide that is used to reliably label the planarian CNS, and quantified the brain-to-body size ratio (<u>Agata et al., 1998</u>; <u>Roberts-Galbraith et al., 2016</u>) to determine if PCP loss affected brain size (Fig. 4B). When normalized to body size at 8 weeks, we found that PCP RNAi animals had significantly larger brains (on average 21% larger than controls, p ≤ 0.01, Fig. 4C), further suggesting that PCP loss leads to neural hyperplasia. To determine if increased brain size resulted from changes in individual subtypes, we investigated the expression of three different neuronal classes that all arise from the same progenitor cell type: GABAergic neurons (*smed-gad)*, octopaminergic neurons (*smed-tbh*), and dopaminergic neurons (*smed-th*) (<u>Cowles et al., 2013</u>). When we visualized populations of *gad^+^*, *tbh^+^*, and *th^+^*neurons with FISH and quantified their density in the brain at 8-weeks following PCP inhibition (Fig. 4D-F), we found that neuronal density of each tested population increased significantly following PCP loss (p ≤ 0.01 for each, Fig. 4G). Specifically, the number of *gad^+^*, *tbh^+^*, and *th^+^* neurons (normalized to body area) increased by 40%, 57%, and 41% respectively, as compared to controls. These data suggest that neural hyperplasia resulting from PCP loss is not lineage-specific and highlight a role for PCP signaling in maintaining cell-type allometry in multiple tissue types.

**Figure 4:**
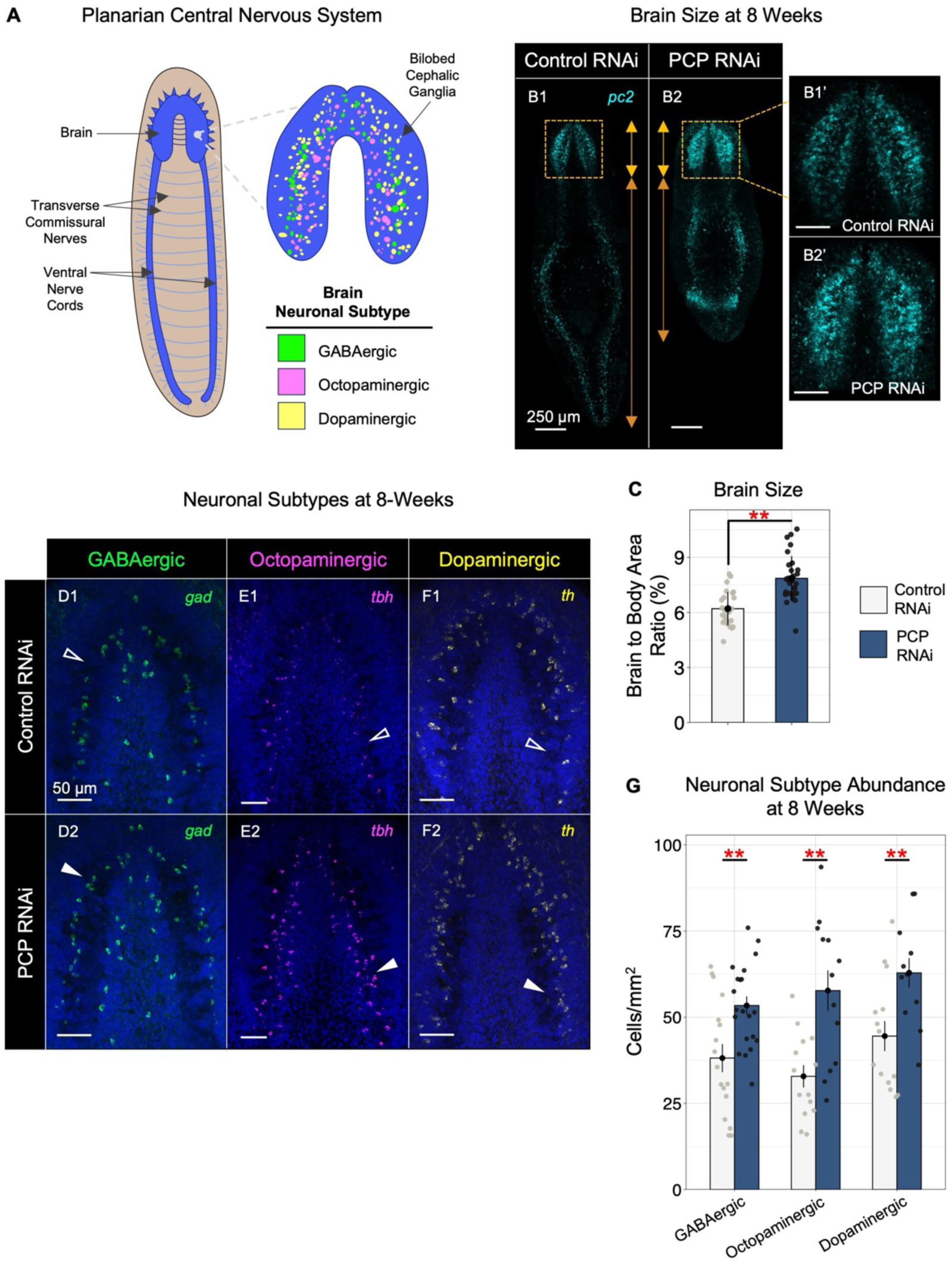
Cell-type allometry is maintained after PCP loss. **(A)** CNS morphology and neuronal subtype distribution in the brain. **(B)** Prohormone convertase-2 (*pc2*) expression at 8 weeks. Orange box: Area shown in B’. Orange arrows: Brain and body lengths. Scale bars in (B): 250 μm. Scale bars in B’: 50 μm. **(C)** Quantification (B); Mean±s.e.m. brain to body area ratio at 8 weeks. n ≥ 27. Two sample *t*-test: ** p≤ 0.01. **(D-F)** Neuronal subtypes (GABAergic (*gad*), octopaminergic (*tbh*), and dopaminergic (*th*) in the brain at 8 weeks. Scale bars for all: 50 µm. White arrowheads = presence of increased neuron numbers. **(G)** Quantification of (D-F); Mean±s.e.m. density of neuronal subtypes in the brain (neurons/mm^2^). n ≥ 12. Two sample *t*-test between Control and PCP RNAi conditions: ** p ≤ 0.01. For all: anterior is up. Ventral view. All experiments were repeated in triplicate.

### PCP Signaling is Required to Limit Progenitor Cell Formation and Maintain tspan-1+ Pluripotent Adult Stem Cell Populations

In addition to body-wide expression in differentiated cell types, *vang-1*, *rock*, and *daam-1* are also expressed in each of the 12 previously described stem cell subtypes (<u>Zeng et al., 2018</u>), including those that give rise to the epidermis, muscle, gut, pharynx, neural tissue, and protonephridia. Interestingly, PCP genes were also enriched in the pluripotent population that is capable of giving rise to all differentiated cell types after lethal irradiation (<u>Zeng et al., 2018</u>). This data, coupled with increases in whole body cell numbers and multi-organ system tissue hyperplasia, led us to hypothesize that PCP signaling may regulate stem cell proliferation and differentiation dynamics during growth termination. To test this hypothesis, we first identified a list of marker genes for several subtypes of planarian stem and progenitor cells from the literature. While all planarian stem and progenitor cells express the PIWI/Argonaute gene *piwi-1* (<u>Reddien, 2005</u>), planarian stem cell populations are highly heterogenous; only a subset of cells are mitotically active and capable of either self-renewal or production of progeny with the capacity to differentiate (Fig. 5A). Whether some populations are more or less pluripotent than others is debated, but *tspan-1* is an established marker of a pluripotent stem cell subpopulation with the capacity to rescue lethally irradiated animals (<u>Zeng et al., 2018</u>). Conversely, progenitor cells express lineage-specific genes and are not mitotically active but do represent a population of precursors to terminally differentiated cells (<u>Pearson, 2022</u>).

**Figure 5:**
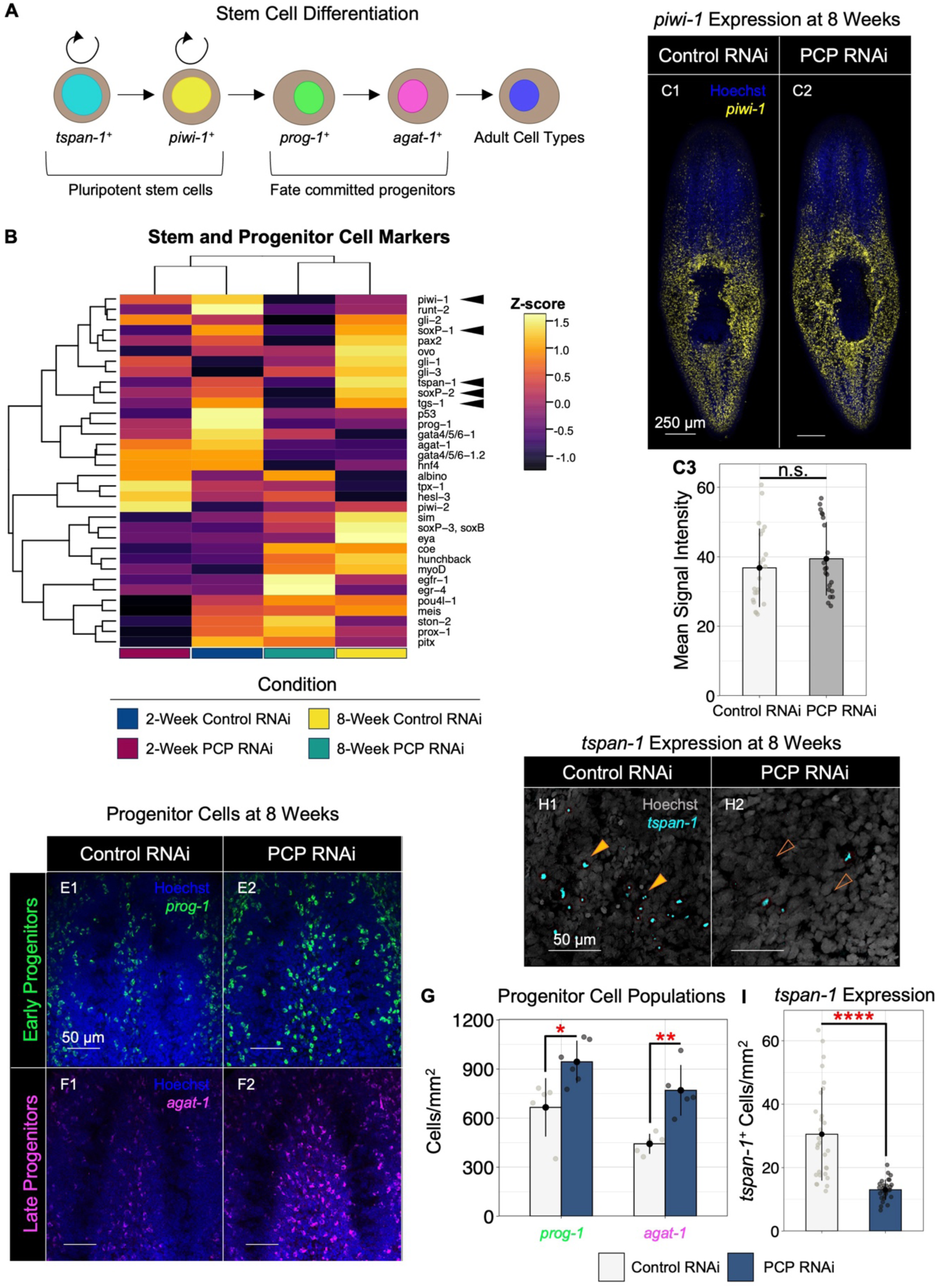
PCP signaling is required to control progenitor cell formation and maintain populations of *tspan-1+* pluripotent stem cells. **(A)** Stages of stem cell differentiation. **(B)** Relative expression (Z-score average FPKM) of known stem and progenitor cell markers in Control and PCP-inhibited animals at 2- and 8-weeks. Markers of pluripotentcy highlighted with black arrowheads. **(C)** Pluripotent stem cells marked by *piwi-1* expression in whole animals at 8 weeks (Blue: nuclei). Scale bars: 250 μm. (C3) Quantification of (C1-2); Mean±s.d. *piwi-1* signal intensity. n ≥ 21. Two sample *t*-test: n.s. **(E-F)** Visualization of early (E, *prog-1*) and late (F, *agat-1*) epidermal progenitor markers. Nuclei: Blue. Scale bars for all: 50 μm. **(G)** Quantification of (E-F); Mean±s.d. early and late progenitor cell density (prog-1^+^ or agat-1^+^ cells/mm^2^). n ≥ 5. Two sample *t*-test: * p < 0.05, ** p ≤ 0.01. **(H)** Visualization of the pluripotent stem cell marker *tspan-1* expression at 8 weeks. Scale bars: 50 µm. Open orange arrowheads: reduction of *tspan-1* expression. **(I)** Quantification of (H); Mean±s.d. *tspan-1^+^* cell density in whole animals following PCP-inhibition. n ≥ 29. Two sample *t*-test: **** p ≤ 0.0001. All experiments were repeated in triplicate.

Here, we compared 34 different planarian stem and progenitor cell markers (<u>King et al., 2024</u>; <u>Molina and Cebria, 2021</u>) and used normal hierarchical clustering to identify expression patterns across conditions in our RNAseq data (Fig. 5B). At 2 weeks, PCP-inhibited animals had increased relative expression of *hesl-3* and *piwi-2* (Fig. 5B) compared to controls at the same timepoint, as well as compared to both conditions at 8-weeks. Both *hesl-3* and *piwi-2* are expressed in dividing stem cells (<u>Cowles et al., 2013</u>; <u>Reddien, 2005</u>); loss of *hesl-3* leads to regeneration defects in the CNS, whereas *piwi-2* is required for regeneration of multiple lineages and maintenance of stem cell populations (<u>Cowles et al., 2013</u>; <u>Reddien, 2005</u>). These results are in line with our previous reports of prolonged stem cell hyperproliferation following PCP inhibition (<u>Beane et al., 2012a</u>). Similarly, we observed increased relative expression of *ston-2*, *prox-1*, and *pitx*, at 8-weeks in PCP-inhibited animals compared to time-matched controls (Fig. 5B). These three genes are all markers of lineage-committed adult stem cells or their progeny, where *ston-2* and *pitx* specify neural fate (<u>Marz et al., 2013</u>; <u>Molinaro and Pearson, 2016</u>), and *prox-1* marks gut-specific progenitors (<u>van Wolfswinkel et al., 2014</u>). Furthermore, we also identified progenitor markers *egr-4* (expressed in many SC subtypes (<u>Zeng et al., 2018</u>)) and *egfr-1* (gut progenitor marker (<u>Barberan et al., 2016</u>)) as having enriched expression in PCP-inhibited animals at 8-weeks (Fig. 5B).

Next, we queried the expression of markers for multilineage and pluripotent stem cells including *tspan-1*, *tgs-1*, *piwi-1*, *soxP-1,* and *soxP-2* (<u>Reddien, 2005</u>; <u>van Wolfswinkel et al., 2014</u>; <u>Zeng et al., 2018</u>). Surprisingly, our data show increased relative expression of *tspan-1*, *tgs-1*, *soxP-1*, and *soxP-2* in 8-week control RNAi animals versus 8-week PCP-inhibited animals, whereas levels of *piwi-1* expression were similar (Fig. 5B). We sought to verify this finding by visualizing expression of *piwi-1* with FISH and quantifying fluorescence intensity as a readout of expression (Fig. 5C). Despite having increased levels of stem cell proliferation at 8 weeks after PCP inhibition (<u>Beane et al., 2012a</u>), we did not detect a significant difference in the expression of *piwi-1* at 8-weeks (p ≥ 0.05, Fig. 5C3), indicating that the effect of PCP loss on stem cell division and differentiation dynamics may be more complex. Although *piwi-1* expression was not affected, we detected increased relative expression levels of multiple progenitor cell markers in our RNA-seq dataset following PCP RNAi at 8 weeks (Fig. 5B). Thus, we wondered whether PCP loss may lead to changes in progenitor cell population numbers (which would also explain the observed multi-organ system hyperplastic phenotypes). Considering PCP-loss led to increased epidermal cell numbers, we visualized the distribution of anterior *prog-1* and *agat-1^+^* epidermal progenitors (<u>Eisenhoffer et al., 2008</u>) with FISH (Fig. 5E-F) and quantified their relative densities. At 8-weeks, PCP RNAi animals had significantly higher densities of both *prog-1^+^* and *agat-1*^+^ cells in the anterior (*prog-1*, 41.8% increase, p ≤ 0.05; *agat-1*, 73.9% increase, p ≤ 0.01, Fig. 5G) as compared to time-matched control RNAi animals.

These data, coupled with the lack of differential *piwi-1* expression (Fig. 5B-C), suggest that stem cell hyperproliferation could result in progenitor cells accumulating at the expense of maintaining pluripotent stem cell populations. Furthermore, this hypothesis is supported by the reduced expression of pluripotent stem cell markers like *tspan-1* and *tgs-1* in PCP-inhibited animals at 8 weeks (Fig. 5B). To test our hypothesis, we visualized *tspan-1* expression and quantified the total number of *tspan-1*^+^ cells per mm^2^ body area in control and PCP-inhibited animals at 8-weeks (Fig. 5H-I). Our data show that loss of PCP signaling leads to significant reductions in whole body numbers of *tspan-1^+^* cells at 8-weeks, (42.7%, p ≤ 0.0001, Fig. 5I), consistent with the observed decrease in markers of pluripotent stem cells in our RNA-seq dataset (Fig. 5B). Collectively, based on our assessment of stem cell differentiation dynamics, hyperproliferation resulting from loss of PCP signaling causes depletion of pluripotent stem cells and accumulation of progenitor and differentiated cells, ultimately leading to body-wide tissue hyperplasia and failure to terminate regenerative growth.

### PCP Signaling is Required to Control Symmetric Versus Asymmetric Division of Stem Cells to Maintain Pluripotency During Tissue Maintenance

Stem cell loss is a hallmark of aging. During normal aging, stem cells become senescent due to defective proteostasis, accumulation of DNA damage, or from remaining in a prolonged state of hyperproliferation that leads to stem cell exhaustion (<u>Huang et al., 2022</u>). Any of these scenarios could explain our observed depletion of *tspan-1^+^* pluripotent stem cells following PCP loss. Another explanation for stem cell loss is an increase in propensity to divide to produce two cells of different fates (instead of dividing to expand pluripotent populations). Symmetric cell division (SCD) produces two identical daughter cells that either maintain stem cell-like properties or become more differentiated progenitors; asymmetric cell division (ACD) generates two cells that are not alike, typically one daughter stem cell and one differentiated progenitor daughter (or two different types of differentiated daughters) (<u>Morrison and Kimble, 2006</u>; <u>Sunchu and Cabernard, 2020</u>). Following sublethal irradiation, planarian stem cells undergo symmetric and asymmetric division during clonal expansion in which approximately 50% of stem cells divide symmetrically (<u>Lei et al., 2016</u>; <u>Raz et al., 2021</u>). These symmetric divisions are characterized by equal inheritance of *piwi-1* mRNA and chromatoid bodies (a stem cell-specific structure; (<u>Kashima et al., 2016</u>), suggesting a proliferative stem cell state. The asymmetrically dividing stem cells produce one *piwi-1*^high^ daughter that remains pluripotent and one *piwi-1*^low^ daughter that will acquire a different cell fate, implying a steady state (<u>Lei et al., 2016</u>; <u>Raz et al., 2021</u>). Recent work shows that pluripotent stem cell populations expand relative to other cell types in smaller animals (<u>Emili et al., 2023</u>), suggesting that periods of tissue maintenance and degrowth promote proliferative SCD. Given that levels of cell death increase during tissue maintenance (<u>Pellettieri et al., 2010</u>), and markers of pluripotency have higher relative expression in control RNAi animals at 8-versus 2-weeks (Fig. 5B), we wondered if loss of pluripotent stem cells following PCP loss may be due to a failure to expand the stem cell pool through symmetric divisions.

Interestingly, many genes associated with SCD versus ACD in other organisms (<u>di Pietro et al., 2016</u>; <u>Lechler and Mapelli, 2021</u>; <u>Lu and Johnston, 2013</u>; <u>Noatynska et al., 2012</u>) appeared as shared DEGs (p ≤ 0.05) in our differential expression analyses (Table S3). We assessed their relative expression (Z-score of average FPKM) in control RNAi and PCP-inhibited animals (Fig. 6A). Although we did not identify any gene-specific expression patterns, normal hierarchical clustering grouped control RNAi animals at 2-weeks with PCP-inhibited animals at 8-weeks, and PCP RNAi animals at 2-weeks with control RNAi animals at 8-weeks (Fig. 6A). Thus, gene expression associated with SCD versus ACD in PCP-inhibited animals more closely matches that observed in 2-week controls, which is interesting considering new tissue formation is sustained in both conditions. These results led us to ask whether PCP signaling may affect the numbers of stem and differentiated cells through regulation of cell division symmetry.

**Figure 6:**
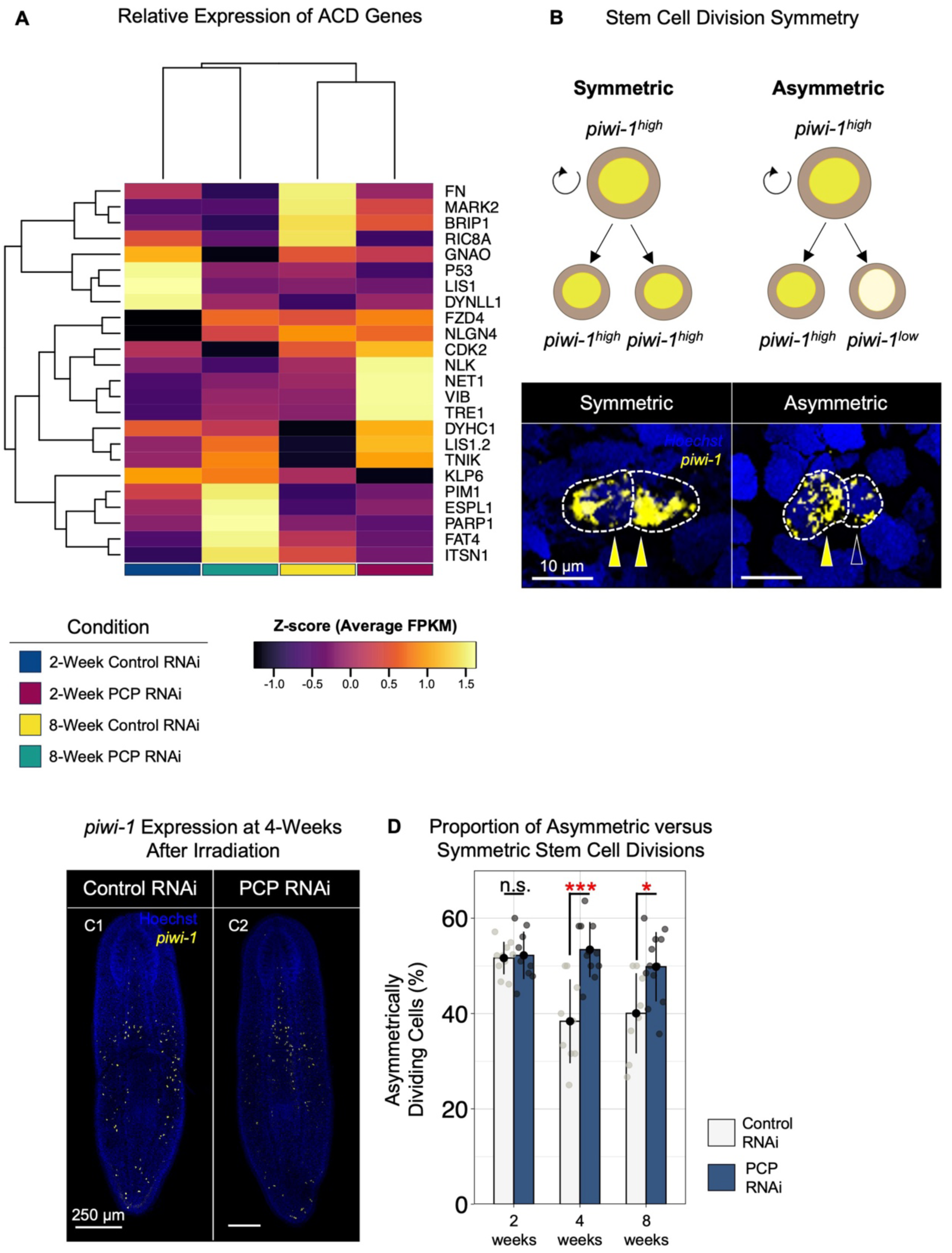
PCP signaling is required to control the symmetric versus asymmetric division of stem cells to maintain pluripotency or produce differentiated daughter cells. **(A)** Relative expression of genes associated with ACD in Control and PCP-inhibited animals at 2- and 8 weeks. Expression values: Z-score of the average FPKM. **(B)** Depiction of *piwi-1* inheritance in planarians stem cells with *in situ* examples of cells undergoing symmetric (left) and asymmetric (right) division. **(C)** Visualization of *piwi-1^+^* clonogenic stem cells in 4-week Control and PCP RNAi animals 5 days post sublethal irradiation (10 Gy). Nuclei: Blue. Scale bars: 250 µm. **(D)** Quantification of (C); Mean±s.d. proportion of *piwi-1^+^* cells undergoing symmetric and asymmetric cell division at 2-, 4-, and 8 weeks. n ≥ 9. Two sample *t*-test between control and PCP-inhibited animals: n.s., *** p ≤ 0.001, * p < 0.05. Two sample *t*-test between Control RNAi animals at 2- and 4-weeks: p = 0.0017. Two sample *t*-test between Control RNAi animals at 2- and 8-weeks: p = 0.0030. All experiments were repeated in triplicate.

To test differences in relative rates of ACD in control and PCP-inhibited animals, we partially depleted the *piwi-1*^+^ stem cell pool using sublethal X-irradiation (as previously described; (<u>Filippova et al., 2023</u>) and allowed stem cell populations to recover for 5 days post-irradiation (Fig. 6B-C). We then assessed ACD versus SCD by visualizing *piwi-1* mRNA distribution in dividing stem cell pairs (<u>Lei et al., 2016</u>; <u>Raz et al., 2021</u>). In this paradigm, symmetrically dividing stem cells are both *piwi-1*^high^ while asymmetrically dividing stem cells produce one *piwi-1*^high^ stem cell and one *piwi-1*^low^ progenitor cell daughter (Fig. 6B). Total numbers of symmetrically and asymmetrically dividing pairs were quantified at 2- , 4-, and 8-weeks (Fig. 6D). The rate of ACD at 2 weeks in control RNAi animals was 51.6%, closely matching previous reports (<u>Lei et al., 2016</u>; <u>Raz et al., 2021</u>). Over time, the rate of ACD was reduced by ∼ 25% in controls at 4-weeks (average rate of ACD: 38.4%, p ≤ 0.01) and at 8-weeks (average rate of ACD: 40.0%, p ≤ 0.01). An increase in SCD rates and pluripotent stem cells over time is consistent with the increase in relative expression of markers of pluripotency at 8 weeks in control RNAi animals (Fig. 5B,I). The rate of ACD in PCP RNAi animals was not significantly different from controls at 2-weeks (average rate of ACD: 52.2%, p ≥ 0.05). However, the rate of ACD in PCP-inhibited animals remained high at both 4 and 8 weeks; 53.4% of stem cell pairs at 4 weeks and 49.8% at 8 weeks underwent ACD (rates of ACD compared to time-matched controls: 2-weeks p ≤ 0.001, 8-weeks p ≤ 0.05) (Fig. 6D). These data indicate that termination of regenerative growth and the initiation of tissue maintenance programs involves a shift in relative rates of ACD and SCD. Specifically, increases in SCD expands pools of pluripotent stem cells, potentially compensating for increased rates of cell loss during homeostasis. Furthermore, PCP signaling is required for a shift from an ACD- to an SCD-dominant cell division program during the termination phase of regeneration, and loss of PCP leads to impaired expansion of the stem cell pool with concurrent depletion of pluripotent populations.

## Discussion

Expanding on our prior work reporting that loss of PCP signaling promotes continued neurogenesis (<u>Beane et al., 2012a</u>), here we demonstrate that PCP signaling is required for termination of body-wide regrowth of gut, excretory, pharyngeal, and epidermal tissues. Through transcriptional profiling of control and PCP-inhibited animals at the end-stages of regeneration and during continued tissue maintenance, we identified differential regulation of planarian stem cell pluripotency genes and show that progenitor cell populations increase in PCP-inhibited animals at the expense of maintaining stem cells. Furthermore, our RNA-seq analyses of control RNAi animals over time provide a resource for scientists interested in other potential growth termination genes or regulators of homeostatic tissue maintenance during periods of degrowth. Finally, we demonstrate that PCP signaling is required to drive symmetric division and pluripotent population expansion at the end of regeneration whereas loss of PCP signaling promotes asymmetric stem cell division and hyperplastic growth.

By combining our transcriptional analysis with publicly available sequencing data (<u>Plass et al., 2018</u>; <u>Rozanski et al., 2019</u>), we found that prospective growth termination DEGs associated with PCP signaling were also differentially expressed upon *hippo*, *yorkie*, or *p53* RNAi (<u>de Sousa et al., 2018</u>; <u>Lin and Pearson, 2014</u>; <u>Lin and Pearson, 2017</u>; <u>Tu et al., 2015</u>). While *p53* was significantly downregulated following PCP-inhibition at 2-weeks (Table S2), we did not detect differential expression of either *hippo* or *yorkie* in any of our data sets. Interestingly, at 8-weeks following PCP loss, we identified the atypical cadherin Fat-4 and injury-induced wound response genes like *egr-4*, *egr-3*, and *hsp70* as significantly upregulated (Table S2) (<u>Owlarn et al., 2017</u>; <u>Wenemoser et al., 2012</u>; <u>Wurtzel et al., 2015</u>). Because prolonged injury signaling can lead to hyperproliferation and expansion of differentiated cell populations (<u>Lin and Pearson, 2014</u>; <u>Lin and Pearson, 2017</u>), growth termination mechanisms likely include transcriptional repression of injury signals. Thus, our data suggest that PCP signaling suppresses growth-inducing transcriptional programs.

While genetic regulators of stem cell proliferation and fate decisions have been studied for decades, biomechanical signals such as shear stress, increasingly recognized as important signals for control of stem cell activities, are less well understood (<u>Vining and Mooney, 2017</u>). Candidates for mediating such mechanosensitive signal transduction in the context of growth termination include Hippo/Yap, Piezo1/p53, Fat/Dachsous/Four-Jointed, and core PCP pathways (<u>Bosveld et al., 2012</u>; <u>Cai et al., 2021</u>; <u>Cheong et al., 2020</u>; <u>Esposito et al., 2022</u>; <u>Peng et al., 2022</u>). These have been thoroughly investigated in hemodynamic regulation of hematopoietic stem cell (HSC) development from the hemogenic endothelium in the aorta-gonad-mesonophros region (<u>Li et al., 2021</u>) along with Notch signaling in the context of HSC development and balancing quiescence with long-term self-renewal (<u>Butko et al., 2016</u>). Moreover, in lung epithelial cells, mechanosignaling was reported to regulate PCP signaling by controlling YAP transduction to the nucleus (<u>Cheong et al., 2020</u>). Our data, which suggests that PCP signaling instructs the intracellular machinery that regulates stem cell proliferation and cell fate decisions during growth suppression, extends significantly from this work and opens up novel research directions. Specifically, more work is needed to elucidate whether core PCP components are mechanosensitive in the context of growth regulation in other biological settings and in other adult stem cell populations, and if so, what the underlying regulatory mechanisms are.

Genes in the *blitzschnell* family has been reported to regulate whole body cell numbers by balancing cell death and stem cell proliferation via the insulin/AKT/mTOR signaling pathway (<u>Pascual-Carreras et al., 2020</u>). Under starvation conditions, *blitzschnell* (*bls*) RNAi results in tissue overgrowth due to a hyperproliferation phenotype in which cell size is reduced and cell numbers increased (<u>Pascual-Carreras et al., 2020</u>). While the hyperplastic phenotype we report here is similar to loss of *blitzschnell,* unlike in our study loss of *blitzschnell* leads to formation of undifferentiated tissue masses (<u>Pascual-Carreras et al., 2020</u>). This difference suggests that growth regulation orchestrated by PCP signaling is mechanistically different from the reported metabolic regulation of cell numbers and body size through *blitzschnell* signaling.

Stem cells prevent exhaustion through regulation of their proliferative state and simultaneously maintain their populations through symmetric versus asymmetric divisions (<u>Obernier et al., 2018</u>; <u>Sunchu and Cabernard, 2020</u>). Our data indicate that stem cells increasingly undergo proliferative symmetric division to expand cell populations as the regeneration permissive environment returns to a state of homeostasis, consistent with recent work (<u>Emili et al., 2023</u>; <u>Gonzalez-Estevez et al., 2012</u>). We further show that loss of PCP signaling leads to a failure to repress the asymmetric stem cell divisions that produce progenitor cells and differentiated tissues (Fig. 6). Consistent with our findings, recent evidence suggests that muscle stem cells can switch between ACD and SCD during periods of regeneration (<u>Evano et al., 2020</u>). Our study extends from this work by suggesting a new role for PCP signaling in regulating this regeneration ’switch’. Future studies are needed to elucidate the specific role of these PCP-mediated genes in control of ACD versus SCD decisions in pluripotent stem cells.

Collectively, our data shed light on the mechanistic role of PCP signaling in regulating growth termination in adult tissue regeneration. We show that core PCP genes are required for repression of ACD at the end stages of regeneration. In addition, our transcriptional profiling of PCP-inhibited animals points to key mechanosignaling pathways as either downstream targets of PCP-signaling or upstream activators of growth termination pathways. Identifying cell-extrinsic cues and cell-intrinsic genetic programs (as well as communication between them) that regulate growth termination programs has important implications for regenerative medicine, where appropriate spatiotemporal repression of new tissue growth is paramount for preventing unwanted hyperproliferation and potential carcinogenesis. Similarly, furthering our understanding of endogenous growth termination mechanisms presents new opportunities to study how these processes are hijacked during tumorigenesis to promote cancer invasion, progression, and metastasis.

## Materials and Methods

### Animal care

*Schmidtea mediterranea* (asexual, clonal line CIW4) were maintained at 18°C in the dark in 0.5 g/liter Instant Ocean salts (in ultrapure water of Type 1). Animals were fed once per week with paste made from antibiotic-free/hormone-free whole calf liver (Creekstone Farms, Arkansas, KS; or C. Roy’s, Yale, Michigan) that was thawed only once before use. Animals were starved 2 weeks before experimentation.

### RNA interference and Amputations

RNAi was performed via feeding of in vitro–synthesized double-stranded RNAi, as previously described (<u>Rouhana et al., 2013</u>). Sequence information for genes used for RNAi is supplied in Table S4. For all experiments, animals (starting size of 2 – 4 mm) were fed RNAi four times over 11 days and amputated into trunk fragments on the 14^th^ day. Control RNAi was double-stranded RNA to Venus-GFP, which is not present in the *S. mediterranea* genome. Animals were amputated via scalpel cuts done under a dissecting microscope on a custom-made cooling Peltier plate, as previously described (<u>Beane et al., 2012a</u>). Regenerating animals were maintained at 20°C in the dark for 2, 4 or 8 weeks. Media was refreshed weekly.

### RNA Extraction, Library Preparation, and RNA Sequencing

A total of n = 6 animals per replicate were used for RNA extractions of Control and PCP RNAi animals at 2-weeks, and n = 10 animals were used per replicate for RNA extractions of Control and PCP RNAi animals at 8-weeks. Numbers of animals per replicate were adjusted so total RNA extracted for each samples was ∼ 5 mg. Each condition was repeated n = 3 times. Tissues were frozen in liquid nitrogen and ground to a fine powder with mortar and pestle and resuspended with lysis buffer from the Qiagen QIAshredder assay kit (Cat. No. 79656). Tissue lysis was performed according to the manufacturer’s protocol.

Total RNA was extracted using the Qiagen RNeasy Plus Micro Kit (Cat. No. 74034) following the manufacturer’s protocol. RNA purity and concentration was assessed using both Invitrogen Qubit RNA High Sensitivity Assay (Cat. No. Q32852) on a Thermo Fisher Qubit Fluorometer and Agilent RNA ScreenTape (No. 5067-5576) on an Agilent TapeStation 2200 according to the manufacturer’s suggested protocols. Samples were stored in nuclease-free water at -80°C prior to library preparation. mRNA was purified from total RNA using poly-T oligo-attached magnetic beads. M-MuLV Reverse Transcriptase (RNase H-, Promega) was used for first-strand cDNA synthesis, while DNA Polymerase I and RNase H (Promega) were used for second strand synthesis. Library fragments were purified with AMPure XP system (Beckman Coulter, Beverly, USA) to select cDNA fragments 370∼420 bp in length. PCR was performed with Phusion High-Fidelity DNA polymerase, Universal PCR primers and Index (X) Primer. Purified PCR products (AMPure XP system) and library quality were assessed using an Agilent Bioanalyzer 2100 system. Index-coded samples were clustered with a cBot Cluster Generation System using TruSeq PE Cluster Kit v3-cBot-HS (Illumia) according to the manufacturer’s protocol. Paired-end sequencing reads of 150 bp were generated from library preparations using an Illumina Novaseq 6000 platform. Raw sequencing reads are freely available under National Library of Medicine BioProject PRJNA1162067.

### Bioinformatics Analysis

An index of the reference genome (*Schmidtea mediterranea* S2F2 genome (<u>Brandl et al., 2016</u>); NCBI Assembly: GCA_0000691995.1) and alignment of clean paired-end reads was performed using Hisat2 v2.0.5(<u>Pertea et al., 2016</u>). Mapped reads were assembled by StringTie (v1.3.3b) (<u>Pertea et al., 2016</u>) using a reference-based approach and gene expression levels (Fragments Per Kilobase of transcript per Million mapped reads, FPKM) were quantified with featureCounts v1.5.0-p3 (<u>Liao et al., 2013</u>). Differential expression analysis was performed using the DESeq2 (1.20.0) (<u>Love et al., 2014</u>) and significance values (p ≤ 0.05) for normalized expression values were adjusted for false discovery rate using the Benjamini and Hochberg’s approach.

Principle component analysis (PCA, Fig. S1) of the whole transcriptome was generated from log_2_FPKM values of each gene for each replicate. Venn diagrams used to visualize similarly differentially expressed genes were produced using Venny 2.1 (<u>Oliveros, 2007-2015</u>). Heatmaps showing shared DEGs between comparisons (Fig. 1) and select stem/progenitor markers and ACD/SCD genes (Fig. 5 and Fig. 6, respectively) were generated using the Z-score of log_2_FPKM averaged across replicates for each condition to show relative changes in gene expression. Dendrograms for column and row organization for all heatmaps were generated by complete, normal hierarchical clustering with the hclust function in R.

### X-Ray Irradiation

Sub-lethally irradiated animals were generated using x-ray irradiation, as previously described (<u>Filippova et al., 2023</u>) with slight modification. Briefly, irradiations were conducted using an X-Rad320 system (Precision X-Ray Irradiation, Set KV: 320.0, Set mA: 12.50) with a total dose of 10 Gy delivered at a rate of 1.3 Gy/min. Animals were placed onto moistened filter paper during treatment and transferred back to planarian salts after treatment. Animals were housed at 20°C in the dark for 5 days prior to fixation (See *in situ hybridization* below for details).

### Whole-mount actin and nuclei labeling and in situ hybridization

Whole mount F-actin labeling was conducted as previously described (<u>Cebrià et al., 1997</u>) with slight modification. Briefly, animals were killed in 7.5% NAC (Sigma A7250) in 1X PBS for 7.5 min and then fixed 4% formaldehyde solution (Supelco, FX0410) in 1X PBS for 20 min. After washing 10 min in 1X PBS, samples were permeabilized/blocked in 1X PBS with 0.3% Triton X and 5% horse serum for 1-2 hours. Samples were washed for 15 min in 1X PBS, then incubated 16-18 hours at 4°C in 0.5 mM FITC-phalloidin (Phalloidin-FITC; Sigma P5282) diluted in 1X PBS. Following incubation, samples were washed 3 times for 10 min with 1X PBS, then counterstained for nuclei (Hoechst 33342 10 mg/mL; ThermoFisher H3570) for 1 hour at RT. Samples were washed 3 times for 10 min in 1X PBS and cleared in 90% glycerol/water solution for several hours before mounting and viewing under a confocal microscope.

Fluorescence *in situ* hybridization (FISH) was carried out as previously described (<u>King and Newmark, 2013</u>), with slight modification; following tyramide signal amplification, samples were washed three times in TNTx and transferred to 1 mg/mL Hoechst 33342 (ThermoFisher H3570) for 16-18 hours at 4°C. Following nuclei counterstaining, samples were washed 6x in TNTx at RT and were then cleared in 90% glycerol/PBS at 4°C. Sequence information for riboprobes is provided in Table S4.

Whole mount colorimetric *in situ* hybridization (WISH) was performed as previously described (<u>Pearson et al., 2009</u>), with slight modifications. Briefly, formaldehyde fixed animals were bleached in 5% formamide and 1.2% H_2_O_2_ bleach for 4 hours under bright light at RT. Tissues were permeabilized with 2 mg/mL proteinase K (Invitrogen 25530049) in PBSTx for 12 minutes at 37°C and then fixed in 4% formaldehyde for 10 minutes. Animals were then incubated at 80°C for 10 minutes to denature endogenous alkaline phosphatase. Following overnight riboprobe hybridization at 56°C and subsequent washes, animals were incubated in a-Digoxigenin-AP fab fragments (Sigma-Aldrich, 11093274910) at a final concentration of 1:300 in blocking solution overnight at 4°C. Following several washes with maleic acid buffer, samples were developed in the dark using SIGMAFAST BCIP/NBT tabs (Sigma-Aldrich, b-5655). For each riboprobe, all samples were developed for the same length of time. Samples were washed and fixed in 4% formaldehyde solution, and ethanol treatment was used to remove background staining prior to clearing in 90% glycerol/PBS at 4°C. Sequence information for riboprobes is provided in Table S4.

### Image collection

Animal whole-body morphology and WISH images were collected using a ZEISS V20 fluorescence stereomicroscope with AxioCam MRc camera and ZEN lite software (ZEISS). For live imaging of whole animals, worms were imaged while fully extended and moving to ensure the absence of tissue bunching.

Images for FISH experiments were collected using a Leica Stellaris 5 upright laser scanning confocal microscope with 3 HyD S detectors (410-850 nm), LIAchroic splitter, and 4 laser diodes (405, 488, 561, and 638 nm) and LasX software. All images were collected with a 0.75 NA 20X Plan Apochromatic air objective, except actin/nuclei labeling (Fig. 3) and *piwi-1^+^*cell division (Fig. 6) which utilized a 1.3 NA 40X Plan Apochromatic oil objective. Unless stated otherwise, images are max intensity projections of stitched z-stacks adjusted linearly for brightness/contrast and improvement of clarity using Adobe Photoshop. Data were neither added nor subtracted; original images are available upon request.

### Flow Cytometric Analyses

All lysates were sorted using a Thermo Fisher Attune NxT flow cytometer. Generation of single cell suspensions: For each replicate, a total of n = 33 animals were used to generate homogenized single cell suspensions, and staining procedures were carried out as in (<u>Peiris et al., 2016</u>). Briefly, animals were diced with a razorblade on a cold plate and transferred to cold calcium-magnesium-free buffer plus 1% bovine serum albumin (CMFB (<u>Reddien, 2005</u>)) pH adjusted to 7.3 and gently rocked for 30 minutes at 4°C to dissociate cells. Prior to staining for flow cytometric analyses, lysates were strained through a 40 mm cell strainer, pelleted by centrifugation at 1500 rpm for 5 minutes (at 4°C), and resuspended in 5 mL ice cold CMFB. Fluorescent labeling: For each experiment, 1 mL of each cell lysate (between 5 x 10^5^ and 1 x 10^6^ cells) was utilized for staining. Cell viability was assessed using trypan blue staining and a hemocytometer. Cells were pelleted again and resuspended in 500 mL of 5 mM DRAQ-5 (Invitrogen eBioscience DRAQ5, 65-0880-96) in CMFB to label nuclei. Lysates were incubated for 10 minutes in the dark at RT before addition of Calcein-AM (Invitrogen, C3099) cell viability marker at a final concentration of 0.15 mg/mL. Cells incubated in the dark for an additional 10 minutes at RT. Cells were pelleted and re-suspended in 100 mL of 1X Annexin-V buffer and 2.5 mL Annexin-V Pacific Blue (Invitrogen, A35136) before incubation at RT for 15 minutes. Finally, lysates were brought to a final volume of 500 mL in 1X Annexin-V buffer before final straining through flow tube caps. Cells were kept in the dark on ice until analysis and the entire volume of lysate was used for analysis. All data analysis was performed using FlowJo software (TreeStar).

### Quantification and statistical analyses

Statistical Analysis: All statistical analyses and graphs were generated using either Microsoft Excel (StatPac, V. 4.0) or RStudio (4.3.0 (2023-04-21), “Already Tomorrow”). Significance was determined with a two-tailed Student’s *t* test with unequal variance for all, except for the following exceptions: Data from the symmetric versus asymmetric cell division quantifications were analyzed by a two-sample *t*-test between percents (two-tailed). Gene Expression *in situ*: Quantification of *piwi-1* expression in whole animals at 8-weeks (Fig. 6): The Adobe Photoshop magnetic lasso tool was used to obtain mean signal intensity values for the whole-body area following FISH. Morphological Analysis of Organ Systems and Stem/Progenitor Cell Numbers: Overall animal size was quantified at 2- and 8-weeks following live animal imaging. Animals were imaged while fully extended and moving to avoid tissue bunching. The magnetic lasso tool in Adobe Photoshop was used to measure animal areas (Fig. 3). The Adobe Photoshop magnetic lasso tool was used to measure the area of the brain and body, and the brain to body size ratio for each animal was expressed as brain size as a percentage of body size (Fig. 4). Similarly, the pharynx to body size ratio utilized the same method, where pharynx area (measured with the magnetic lasso tool) was normalized to body area and expressed as a percentage (Fig. S2). The number of *gad*, *tbh*, and *th^+^*neurons were quantified in the area between cephalic ganglia and normalized to animal body size, where the number of neurons is expressed as the number of positive cells/mm^2^ body area (Fig. 4). Similarly, protonephridia or secondary intestinal branches were counted and normalized to animal body area and are expressed as protonephridia/mm^2^ or secondary branches/mm^2^, respectively (Fig. 2). For analysis of the dimensions of individual secondary intestinal branches: The Adobe Photoshop ruler tool was used to measure either the length of each branch from the base (where it attaches to the primary intestinal branch) or the width of the branch at the base. The length of the branch was normalized to the width of the animal, whereas width measurements were normalized to the primary branch length (Fig. 2). For quantification of epidermal cell density: The total number of epidermal nuclei were counted in a 50 mm x 50 mm area in the prepharyngeal region using the threshold feature in FIJI/ImageJ (Fig. 5). Finally, the numbers of *prog-1* and *agat-1*^+^ cells were counted in the area between cephalic ganglia following FISH, and total counts were normalized to the area between both lobes. Cell counts are expressed as *prog-1* or *agat-1^+^* cells/mm^2^ (Fig. 5). *tspan-1^+^* cells were visualized with FISH; the number of *tspan-1^+^* cells was quantified and normalized to body area (measured with the magnetic lasso tool). Cell counts are expressed as *tspan-1^+^* cells/mm^2^ (Fig. 5). Symmetric versus Asymmetric Cell Division Analysis: Proportions of symmetric versus asymmetric cell divisions in *piwi-1^+^* cells were quantified in irradiated Control and PCP RNAi animals at 5 dpi as follows: Max intensity projections were generated from Hoechst and *piwi-1* channels from 35 um Z-stack images (20 slices, 1.85 um/slice) of whole animals. The number of symmetric and asymmetric cell divisions was quantified in the visible area, where the proportion of asymmetric cell divisions is expressed as a percent of the total number of cell divisions as in (<u>Lei et al., 2016</u>; <u>Raz et al., 2021</u>) (Fig. 6).

See *Bioinformatics Analysis* and *Flow Cytometric Analyses* for relevant quantification information.

### Reagent Requests

Requests for any reagents used in this study can be made to the corresponding author upon reasonable request.

## Acknowledgements

This work was supported by National Science Foundation (#1652312) and National Institutes of Health (#1R15GM150073-01) grants to W.S.B and Western Michigan University Dissertation Completion Fellowship to S.J.H. The authors would also like to thank Michael Clemente, Flow Cytometry and Imaging Core Manager, and Alissa Cousino, Core Laboratory Manager, at WMU Homer Stryker M.D. School of Medicine for aiding with flow cytometric analyses and animal irradiations, respectively. Data from this publication has been previously described in a publicly available dissertation (<u>Hack, 2024</u>).

## Author contributions

This work was supervised by W.S.B., who supplied materials and reagents. S.J.H conceived of the initial ideas and wrote the initial manuscript. S.J.H. analyzed all the data, with contributions from W.S.B. S.J.H. performed all the assays. S.J.H and W.S.B. prepared the figures. W.S.B. and S.J.H. edited the draft and final versions of the manuscript.

## Data availability

All relevant data supporting the key findings of this study are available within the article, supplemental information, public repositories, or from the corresponding author upon reasonable request.

## References

Adler, C. E., Seidel, C. W., McKinney, S. A. and Sánchez Alvarado, A. (2014). Selective amputation of the pharynx identifies a FoxA-dependent regeneration program in planaria. Elife 3, e02238.

Agata, K., Soejima, Y., Kato, K., Kobayashi, C., Umesono, Y. and Watanabe, K. (1998). Structure of the planarian central nervous system (CNS) revealed by neuronal cell markers. Zoolog Sci 15, 433–440.

Almuedo-Castillo, M., Salo, E. and Adell, T. (2011). Dishevelled is essential for neural connectivity and planar cell polarity in planarians. Proc Natl Acad Sci U S A 108, 2813–2818.

Barberan, S., Fraguas, S. and Cebria, F. (2016). The EGFR signaling pathway controls gut progenitor differentiation during planarian regeneration and homeostasis. Development 143, 2089–2102.

Barghouth, P. G., Thiruvalluvan, M., LeGro, M. and Oviedo, N. J. (2019). DNA damage and tissue repair: What we can learn from planaria. Semin Cell Dev Biol 87, 145–159.

Beane, W. S., Tseng, A.-S., Morokuma, J., Lemire, J. M. and Levin, M. (2012a). Inhibition of Planar Cell Polarity Extends Neural Growth During Regeneration, Homeostasis, and Development. Stem Cells and Development 21, 2085–2094.

Beane, W. S., Tseng, A. S., Morokuma, J., Lemire, J. M. and Levin, M. (2012b). Inhibition of planar cell polarity extends neural growth during regeneration, homeostasis, and development. Stem Cells Dev 21, 2085–2094.

Bosveld, F., Bonnet, I., Guirao, B., Tlili, S., Wang, Z., Petitalot, A., Marchand, R., Bardet, P. L., Marcq, P., Graner, F. and Bellaïche, Y. (2012). Mechanical control of morphogenesis by Fat/Dachsous/Four-jointed planar cell polarity pathway. Science 336, 724–727.

Brandl, H., Moon, H., Vila-Farré, M., Liu, S. Y., Henry, I. and Rink, J. C. (2016). PlanMine--a mineable resource of planarian biology and biodiversity. Nucleic Acids Res 44, D764–773.

Butko, E., Pouget, C. and Traver, D. (2016). Complex regulation of HSC emergence by the Notch signaling pathway. Dev Biol 409, 129–138.

Butler, M. T. and Wallingford, J. B. (2017). Planar cell polarity in development and disease. Nat Rev Mol Cell Biol 18, 375–388.

Cai, X., Wang, K. C. and Meng, Z. (2021). Mechanoregulation of YAP and TAZ in Cellular Homeostasis and Disease Progression. Front Cell Dev Biol 9, 673599.

Cebrià, F., Vispo, M., Newmark, P., Bueno, D. and Romero, R. (1997). Myocyte differentiation and body wall muscle regeneration in the planarian Girardia tigrina. Dev. Genes Evol. 207, 306–316.

Chavali, M., Klingener, M., Kokkosis, A. G., Garkun, Y., Felong, S., Maffei, A. and Aguirre, A. (2018). Non-canonical Wnt signaling regulates neural stem cell quiescence during homeostasis and after demyelination. Nat Commun 9, 36.

Chen, W. S., Antic, D., Matis, M., Logan, C. Y., Povelones, M., Anderson, G. A., Nusse, R. and Axelrod, J. D. (2008). Asymmetric homotypic interactions of the atypical cadherin flamingo mediate intercellular polarity signaling. Cell 133, 1093–1105.

Cheong, S. S., Akram, K. M., Matellan, C., Kim, S. Y., Gaboriau, D. C. A., Hind, M., Del Río Hernández, A. E., Griffiths, M. and Dean, C. H. (2020). The Planar Polarity Component VANGL2 Is a Key Regulator of Mechanosignaling. Front Cell Dev Biol 8, 577201.

Cowles, M. W., Brown, D. D., Nisperos, S. V., Stanley, B. N., Pearson, B. J. and Zayas, R. M. (2013). Genome-wide analysis of the bHLH gene family in planarians identifies factors required for adult neurogenesis and neuronal regeneration. Development 140, 4691–4702.

Cowles, M. W., Omuro, K. C., Stanley, B. N., Quintanilla, C. G. and Zayas, R. M. (2014). COE loss-of-function analysis reveals a genetic program underlying maintenance and regeneration of the nervous system in planarians. PLoS Genet 10, e1004746.

Csibi, A. and Blenis, J. (2012). Hippo-YAP and mTOR pathways collaborate to regulate organ size. Nat Cell Biol 14, 1244–1245.

de Sousa, N., Rodríguez-Esteban, G., Rojo-Laguna, J. I., Saló, E. and Adell, T. (2018). Hippo signaling controls cell cycle and restricts cell plasticity in planarians. PLoS Biol 16, e2002399.

Devenport, D., Oristian, D., Heller, E. and Fuchs, E. (2011). Mitotic internalization of planar cell polarity proteins preserves tissue polarity. Nat Cell Biol 13, 893–902.

di Pietro, F., Echard, A. and Morin, X. (2016). Regulation of mitotic spindle orientation: an integrated view. EMBO Rep 17, 1106–1130.

Donthamsetty, S., Bhave, V. S., Mars, W. M., Bowen, W. C., Orr, A., Haynes, M. M., Wu, C. and Michalopoulos, G. K. (2013). Role of PINCH and its partner tumor suppressor Rsu-1 in regulating liver size and tumorigenesis. PLoS One 8, e74625.

Eisenhoffer, G. T., Kang, H. and Sanchez Alvarado, A. (2008). Molecular analysis of stem cells and their descendants during cell turnover and regeneration in the planarian Schmidtea mediterranea. Cell Stem Cell 3, 327–339.

Emili, E., Pérez-Posada, A., Christodoulou, M. D. and Solana, J. (2023). Allometry of cell types in planarians by single cell transcriptomics. bioRxiv, 2023.2011.2001.565140.

Esposito, D., Pant, I., Shen, Y., Qiao, R. F., Yang, X., Bai, Y., Jin, J., Poulikakos, P. I. and Aaronson, S. A. (2022). ROCK1 mechano-signaling dependency of human malignancies driven by TEAD/YAP activation. Nat Commun 13, 703.

Evano, B., Khalilian, S., Le Carrou, G., Almouzni, G. and Tajbakhsh, S. (2020). Dynamics of Asymmetric and Symmetric Divisions of Muscle Stem Cells In Vivo and on Artificial Niches. Cell Rep 30, 3195–3206.e3197.

Filippova, K. O., Ermakov, A. M., Popov, A. L., Ermakova, O. N., Blagodatsky, A. S., Chukavin, N. N., Shcherbakov, A. B., Baranchikov, A. E. and Ivanov, V. K. (2023). Mitogen-like Cerium-Based Nanoparticles Protect Schmidtea mediterranea against Severe Doses of X-rays. Int J Mol Sci 24.

Fincher, C. T., Wurtzel, O., de Hoog, T., Kravarik, K. M. and Reddien, P. W. (2018). Cell type transcriptome atlas for the planarian Schmidtea mediterranea. Science 360.

Flores, N. M., Oviedo, N. J. and Sage, J. (2016). Essential role for the planarian intestinal GATA transcription factor in stem cells and regeneration. Dev Biol 418, 179–188.

Forsthoefel, D. J., Cejda, N. I., Khan, U. W. and Newmark, P. A. (2020). Cell-type diversity and regionalized gene expression in the planarian intestine. Elife 9.

Forsthoefel, D. J., James, N. P., Escobar, D. J., Stary, J. M., Vieira, A. P., Waters, F. A. and Newmark, P. A. (2012). An RNAi screen reveals intestinal regulators of branching morphogenesis, differentiation, and stem cell proliferation in planarians. Dev Cell 23, 691–704.

Forsthoefel, D. J., Park, A. E. and Newmark, P. A. (2011). Stem cell-based growth, regeneration, and remodeling of the planarian intestine. Dev Biol 356, 445–459.

Gallai, M., Sebestyén, A., Nagy, P., Kovalszky, I., Onody, T. and Thorgeirsson, S. S. (1996). Proteoglycan gene expression in rat liver after partial hepatectomy. Biochem Biophys Res Commun 228, 690–694.

González-Estévez, C. (2009). Autophagy meets planarians. Autophagy 5, 290–297.

Gonzalez-Estevez, C., Felix, D. A., Rodriguez-Esteban, G. and Aboobaker, A. A. (2012). Decreased neoblast progeny and increased cell death during starvation-induced planarian degrowth. Int J Dev Biol 56, 83–91.

Grohme, M. A., Schloissnig, S., Rozanski, A., Pippel, M., Young, G. R., Winkler, S., Brandl, H., Henry, I., Dahl, A., Powell, S., et al. (2018). The genome of Schmidtea mediterranea and the evolution of core cellular mechanisms. Nature 554, 56–61.

Hack, S. J. (2024). Cell Signaling Mechanisms in the Initiation, Propagation, and Termination of Adult Tissue Regeneration. In *Department of Biological Sciences*: Western Michigan University.

Hanh Thi-Kim, V., Sarah, M., Michael, K., Corinna, B., Cyril, B., Juliette, A., Eugene Wimberly, M., Lutz, B. and Jochen Christian, R. (2019). Dynamic Polarization of the Multiciliated Planarian Epidermis between Body Plan Landmarks. Developmental Cell 51, 526–542.e526.

Heck, B. W. and Devenport, D. (2017). Trans-endocytosis of Planar Cell Polarity Complexes during Cell Division. Curr Biol 27, 3725–3733.e3724.

Henderson, J. M., Nisperos, S. V., Weeks, J., Ghulam, M., Marín, I. and Zayas, R. M. (2015). Identification of HECT E3 ubiquitin ligase family genes involved in stem cell regulation and regeneration in planarians. Dev Biol 404, 21–34.

Huang, W., Hickson, L. J., Eirin, A., Kirkland, J. L. and Lerman, L. O. (2022). Cellular senescence: the good, the bad and the unknown. Nat Rev Nephrol 18, 611–627.

Ju, R., Cirone, P., Lin, S., Griesbach, H., Slusarski, D. C. and Crews, C. M. (2010). Activation of the planar cell polarity formin DAAM1 leads to inhibition of endothelial cell proliferation, migration, and angiogenesis. Proc Natl Acad Sci U S A 107, 6906–6911.

Kashima, M., Kumagai, N., Agata, K. and Shibata, N. (2016). Heterogeneity of chromatoid bodies in adult pluripotent stem cells of planarian Dugesia japonica. Dev Growth Differ 58, 225–237.

Kim, S. K., Shindo, A., Park, T. J., Oh, E. C., Ghosh, S., Gray, R. S., Lewis, R. A., Johnson, C. A., Attie-Bittach, T., Katsanis, N. and Wallingford, J. B. (2010). Planar cell polarity acts through septins to control collective cell movement and ciliogenesis. Science 329, 1337–1340.

King, H. O., Owusu-Boaitey, K. E., Fincher, C. T. and Reddien, P. W. (2024). A transcription factor atlas of stem cell fate in planarians. Cell Rep 43, 113843.

King, R. S. and Newmark, P. A. (2013). In situ hybridization protocol for enhanced detection of gene expression in the planarian Schmidtea mediterranea. BMC developmental biology 13, 8.

Lapan, S. W. and Reddien, P. W. (2011). dlx and sp6-9 Control optic cup regeneration in a prototypic eye. PLoS Genet 7, e1002226.

Lapan, S. W. and Reddien, P. W. (2012). Transcriptome analysis of the planarian eye identifies ovo as a specific regulator of eye regeneration. Cell Rep 2, 294–307.

Lechler, T. and Mapelli, M. (2021). Spindle positioning and its impact on vertebrate tissue architecture and cell fate. Nat Rev Mol Cell Biol 22, 691–708.

Lei, K., Thi-Kim Vu, H., Mohan, R. D., McKinney, S. A., Seidel, C. W., Alexander, R., Gotting, K., Workman, J. L. and Sánchez Alvarado, A. (2016). Egf Signaling Directs Neoblast Repopulation by Regulating Asymmetric Cell Division in Planarians. Dev Cell 38, 413–429.

Li, H., Luo, Q., Shan, W., Cai, S., Tie, R., Xu, Y., Lin, Y., Qian, P. and Huang, H. (2021). Biomechanical cues as master regulators of hematopoietic stem cell fate. Cell Mol Life Sci 78, 5881–5902.

Liao, Y., Smyth, G. K. and Shi, W. (2013). featureCounts: an efficient general purpose program for assigning sequence reads to genomic features. Bioinformatics 30, 923–930.

Lin, A. Y. and Pearson, B. J. (2014). Planarian yorkie/YAP functions to integrate adult stem cell proliferation, organ homeostasis and maintenance of axial patterning. Development 141, 1197–1208.

Lin, A. Y. T. and Pearson, B. J. (2017). Yorkie is required to restrict the injury responses in planarians. PLoS Genet 13, e1006874.

Love, M. I., Huber, W. and Anders, S. (2014). Moderated estimation of fold change and dispersion for RNA-seq data with DESeq2. Genome Biol 15, 550.

Lu, M. S. and Johnston, C. A. (2013). Molecular pathways regulating mitotic spindle orientation in animal cells. Development 140, 1843–1856.

Luxenburg, C., Heller, E., Pasolli, H. A., Chai, S., Nikolova, M., Stokes, N. and Fuchs, E. (2015). Wdr1-mediated cell shape dynamics and cortical tension are essential for epidermal planar cell polarity. Nat Cell Biol 17, 592–604.

Marinari, E., Mehonic, A., Curran, S., Gale, J., Duke, T. and Baum, B. (2012). Live-cell delamination counterbalances epithelial growth to limit tissue overcrowding. Nature 484, 542–545.

Marz, M., Seebeck, F. and Bartscherer, K. (2013). A Pitx transcription factor controls the establishment and maintenance of the serotonergic lineage in planarians. Development 140, 4499–4509.

Michalopoulos, G. K. and Bhushan, B. (2021). Liver regeneration: biological and pathological mechanisms and implications. Nat Rev Gastroenterol Hepatol 18, 40–55.

Miyamoto, M., Hattori, M., Hosoda, K., Sawamoto, M., Motoishi, M., Hayashi, T., Inoue, T. and Umesono, Y. (2020). The pharyngeal nervous system orchestrates feeding behavior in planarians. Sci Adv 6, eaaz0882.

Molina, M. D. and Cebria, F. (2021). Decoding Stem Cells: An Overview on Planarian Stem Cell Heterogeneity and Lineage Progression. Biomolecules 11.

Molinaro, A. M. and Pearson, B. J. (2016). In silico lineage tracing through single cell transcriptomics identifies a neural stem cell population in planarians. Genome Biol 17, 87.

Morrison, S. J. and Kimble, J. (2006). Asymmetric and symmetric stem-cell divisions in development and cancer. Nature 441, 1068–1074.

Noatynska, A., Gotta, M. and Meraldi, P. (2012). Mitotic spindle (DIS)orientation and DISease: cause or consequence? J Cell Biol 199, 1025–1035.

Obernier, K., Cebrian-Silla, A., Thomson, M., Parraguez, J. I., Anderson, R., Guinto, C., Rodas Rodriguez, J., Garcia-Verdugo, J. M. and Alvarez-Buylla, A. (2018). Adult Neurogenesis Is Sustained by Symmetric Self-Renewal and Differentiation. Cell Stem Cell 22, 221–234.e228.

Oliveros, J. C. (2007-2015). Venny. An interactive tool for comparing lists with Venn’s diagrams.

Oozeer, F., Yates, L. L., Dean, C. and Formstone, C. J. (2017). A role for core planar polarity proteins in cell contact-mediated orientation of planar cell division across the mammalian embryonic skin. Sci Rep 7, 1880.

Owlarn, S., Klenner, F., Schmidt, D., Rabert, F., Tomasso, A., Reuter, H., Mulaw, M. A., Moritz, S., Gentile, L., Weidinger, G. and Bartscherer, K. (2017). Generic wound signals initiate regeneration in missing-tissue contexts. Nat Commun 8, 2282.

Pascual-Carreras, E., Marin-Barba, M., Herrera-Ubeda, C., Font-Martin, D., Eckelt, K., de Sousa, N., Garcia-Fernandez, J., Salo, E. and Adell, T. (2020). Planarian cell number depends on blitzschnell, a novel gene family that balances cell proliferation and cell death. Development 147.

Pearson, B. J. (2022). Finding the potency in planarians. Commun Biol 5, 970.

Pearson, B. J., Eisenhoffer, G. T., Gurley, K. A., Rink, J. C., Miller, D. E. and Sanchez Alvarado, A. (2009). Formaldehyde-based whole-mount in situ hybridization method for planarians. Dev Dyn 238, 443–450.

Peiris, T. H., Garcia-Ojeda, M. E. and Oviedo, N. J. (2016). Alternative flow cytometry strategies to analyze stem cells and cell death in planarians. Regeneration (Oxf) 3, 123–135.

Pellettieri, J., Fitzgerald, P., Watanabe, S., Mancuso, J., Green, D. R. and Sanchez Alvarado, A. (2010). Cell death and tissue remodeling in planarian regeneration. Dev Biol 338, 76–85.

Peng, Y., Du, J., Günther, S., Guo, X., Wang, S., Schneider, A., Zhu, L. and Braun, T. (2022). Mechano-signaling via Piezo1 prevents activation and p53-mediated senescence of muscle stem cells. Redox Biol 52, 102309.

Pertea, M., Kim, D., Pertea, G. M., Leek, J. T. and Salzberg, S. L. (2016). Transcript-level expression analysis of RNA-seq experiments with HISAT, StringTie and Ballgown. Nat Protoc 11, 1650–1667.

Plass, M., Solana, J., Wolf, F. A., Ayoub, S., Misios, A., Glazar, P., Obermayer, B., Theis, F. J., Kocks, C. and Rajewsky, N. (2018). Cell type atlas and lineage tree of a whole complex animal by single-cell transcriptomics. Science 360.

Raz, A. A., Wurtzel, O. and Reddien, P. W. (2021). Planarian stem cells specify fate yet retain potency during the cell cycle. Cell Stem Cell 28, 1307–1322.e1305.

Reddien, P. W. (2018). The Cellular and Molecular Basis for Planarian Regeneration. Cell 175, 327–345.

Reddien, P. W. O., N. J.; Jennings, J. R.; Jenkin, J. C.; Sanchez Alvarado, A. (2005). SMEDWI-2 Is a PIWI-Like Protein That Regulates Planarian Stem Cells. Science 310, 1327–1330.

Roberts-Galbraith, R. H., Brubacher, J. L. and Newmark, P. A. (2016). A functional genomics screen in planarians reveals regulators of whole-brain regeneration. Elife 5.

Rodrigo Albors, A., Tazaki, A., Rost, F., Nowoshilow, S., Chara, O. and Tanaka, E. M. (2015). Planar cell polarity-mediated induction of neural stem cell expansion during axolotl spinal cord regeneration. Elife 4, e10230.

Ross, K. G., Currie, K. W., Pearson, B. J. and Zayas, R. M. (2017). Nervous system development and regeneration in freshwater planarians. Wiley Interdiscip Rev Dev Biol 6.

Rouhana, L., Weiss, J. A., Forsthoefel, D. J., Lee, H., King, R. S., Inoue, T., Shibata, N., Agata, K. and Newmark, P. A. (2013). RNA interference by feeding in vitro synthesized double-stranded RNA to planarians: methodology and dynamics. Developmental dynamics : an official publication of the American Association of Anatomists 242, 718–730.

Rozanski, A., Moon, H., Brandl, H., Martín-Durán, J. M., Grohme, M. A., Hüttner, K., Bartscherer, K., Henry, I. and Rink, J. C. (2019). PlanMine 3.0-improvements to a mineable resource of flatworm biology and biodiversity. Nucleic Acids Res 47, D812–d820.

Rudolph, K. L., Trautwein, C., Kubicka, S., Rakemann, T., Bahr, M. J., Sedlaczek, N., Schuppan, D. and Manns, M. P. (1999). Differential regulation of extracellular matrix synthesis during liver regeneration after partial hepatectomy in rats. Hepatology 30, 1159–1166.

Ségalen, M., Johnston, C. A., Martin, C. A., Dumortier, J. G., Prehoda, K. E., David, N. B., Doe, C. Q. and Bellaïche, Y. (2010). The Fz-Dsh planar cell polarity pathway induces oriented cell division via Mud/NuMA in Drosophila and zebrafish. Dev Cell 19, 740–752.

Shindo, A. and Wallingford, J. B. (2014). PCP and septins compartmentalize cortical actomyosin to direct collective cell movement. Science 343, 649–652.

Shrestha, R., Little, K. A., Tamayo, J. V., Li, W., Perlman, D. H. and Devenport, D. (2015). Mitotic Control of Planar Cell Polarity by Polo-like Kinase 1. Dev Cell 33, 522–534.

Struhl, G., Casal, J. and Lawrence, P. A. (2012). Dissecting the molecular bridges that mediate the function of Frizzled in planar cell polarity. Development 139, 3665–3674.

Strutt, H., Warrington, S. J. and Strutt, D. (2011). Dynamics of core planar polarity protein turnover and stable assembly into discrete membrane subdomains. Dev Cell 20, 511–525.

Sunchu, B. and Cabernard, C. (2020). Principles and mechanisms of asymmetric cell division. Development 147.

Takeda, H., Nishimura, K. and Agata, K. (2009). Planarians maintain a constant ratio of different cell types during changes in body size by using the stem cell system. Zoolog Sci 26, 805–813.

Takemura, M. and Nakato, H. (2017). Drosophila Sulf1 is required for the termination of intestinal stem cell division during regeneration. J Cell Sci 130, 332–343.

Thi-Kim Vu, H., Rink, J. C., McKinney, S. A., McClain, M., Lakshmanaperumal, N., Alexander, R. and Sánchez Alvarado, A. (2015). Stem cells and fluid flow drive cyst formation in an invertebrate excretory organ. Elife 4.

Tu, K. C., Cheng, L. C., H, T. K. V., Lange, J. J., McKinney, S. A., Seidel, C. W. and Sánchez Alvarado, A. (2015). Egr-5 is a post-mitotic regulator of planarian epidermal differentiation. Elife 4, e10501.

Tumaneng, K., Russell, R. C. and Guan, K. L. (2012). Organ size control by Hippo and TOR pathways. Curr Biol 22, R368–379.

Usui, T., Shima, Y., Shimada, Y., Hirano, S., Burgess, R. W., Schwarz, T. L., Takeichi, M. and Uemura, T. (1999). Flamingo, a seven-pass transmembrane cadherin, regulates planar cell polarity under the control of Frizzled. Cell 98, 585–595.

van Wolfswinkel, J. C., Wagner, D. E. and Reddien, P. W. (2014). Single-cell analysis reveals functionally distinct classes within the planarian stem cell compartment. Cell Stem Cell 15, 326–339.

Vining, K. H. and Mooney, D. J. (2017). Mechanical forces direct stem cell behaviour in development and regeneration. Nat Rev Mol Cell Biol 18, 728–742.

Vladar, E. K., Bayly, R. D., Sangoram, A. M., Scott, M. P. and Axelrod, J. D. (2012). Microtubules enable the planar cell polarity of airway cilia. Curr Biol 22, 2203–2212.

Wenemoser, D., Lapan, S. W., Wilkinson, A. W., Bell, G. W. and Reddien, P. W. (2012). A molecular wound response program associated with regeneration initiation in planarians. Genes Dev 26, 988–1002.

Winter, C. G., Wang, B., Ballew, A., Royou, A., Karess, R., Axelrod, J. D. and Luo, L. (2001). Drosophila Rho-associated kinase (Drok) links Frizzled-mediated planar cell polarity signaling to the actin cytoskeleton. Cell 105, 81–91.

Wurtzel, O., Cote, L. E., Poirier, A., Satija, R., Regev, A. and Reddien, P. W. (2015). A Generic and Cell-Type-Specific Wound Response Precedes Regeneration in Planarians. Dev Cell 35, 632–645.

Yang, X., Qian, X., Ma, R., Wang, X., Yang, J., Luo, W., Chen, P., Chi, F. and Ren, D. (2017). Establishment of planar cell polarity is coupled to regional cell cycle exit and cell differentiation in the mouse utricle. Sci Rep 7, 43021.

Yu, F. X., Zhao, B. and Guan, K. L. (2015). Hippo Pathway in Organ Size Control, Tissue Homeostasis, and Cancer. Cell 163, 811–828.

Zeng, A., Li, H., Guo, L., Gao, X., McKinney, S., Wang, Y., Yu, Z., Park, J., Semerad, C., Ross, E., et al. (2018). Prospectively Isolated Tetraspanin(+) Neoblasts Are Adult Pluripotent Stem Cells Underlying Planaria Regeneration. Cell 173, 1593–1608 e1520.

