## Supplementary Material for "Planar cell polarity signaling controls cell division symmetry to promote termination of adult tissue regeneration"

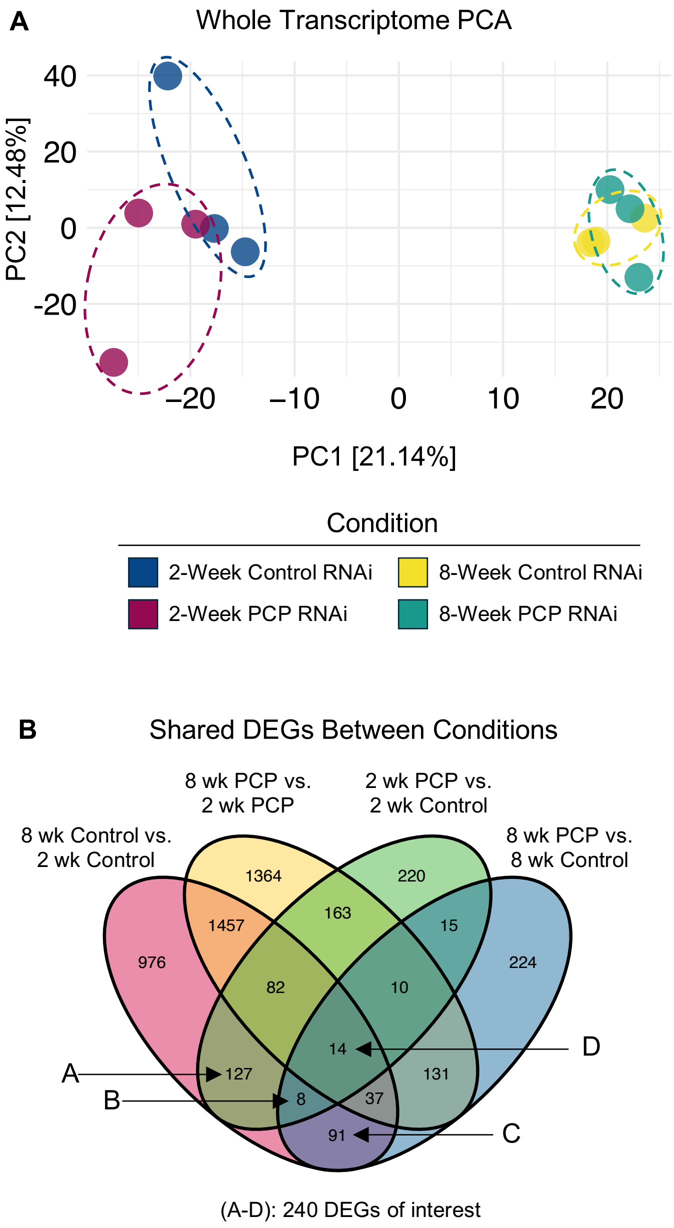


**Figure S1: Transcriptional profiling of PCP-inhibited animals. (A)** Whole transcriptome principal component analysis (PCA) for all replicates of RNA-seq experiment (Fig. 1). Dashed lines: replicates grouped by condition. **(B)** Identification of shared genes between DEG comparisons. Comparison groups containing genes of interest (Fig. 1F) are labeled A-D.

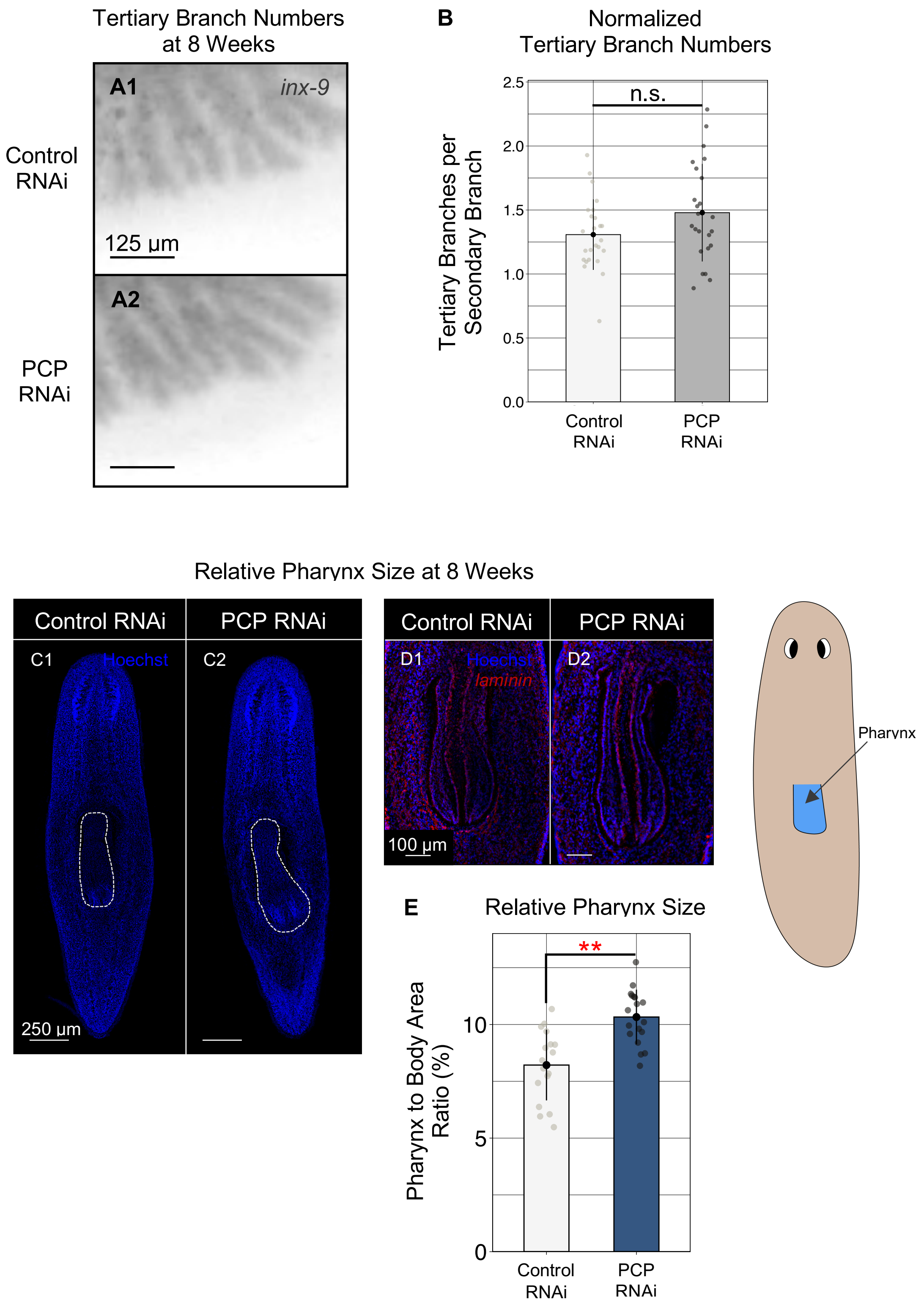


**Fig. S2: Termination of pharynx regeneration requires PCP signaling. (A)** Secondary and tertiary intestinal branches visualized by *inx-9* expression at 8 weeks. Scale bars: 125 µm. **(B)** Quantification of (A); Mean±s.d. tertiary branch numbers at 8 weeks. n ≥ 24. Two sample *t*-test: n.s. **(C-D)** Pharynx morphology at 8 weeks following PCP inhibition. Blue: Nuclei. Red: *laminin* expression. Pharynx outlined in white in (C) is shown in (D). Scales bars: 250 µm (C) and 100 µm (D). Anterior is up. **(E)** Quantification of (C-D); Mean±s.d. pharynx size expressed as pharynx to body area ratio (%). n ≥ 17. Two sample *t*-test: ** p ≤ 0.01. All experiments were repeated in triplicate.
